# Actin cytoskeletal deregulation, caused by RhoGEF2 overexpression, induces cell competition dependent on Ptp10D, Crumbs, and the Hippo signaling pathway

**DOI:** 10.1101/2025.04.26.650750

**Authors:** Natasha Fahey-Lozano, Marta Portela, John E. La Marca, Helena E. Richardson

## Abstract

In *Drosophila* larval epithelial tissues, cells containing mutations in the apico-basal polarity proteins, Scrib, Dlg or Lgl, are eliminated by cell competition when surrounded by *wild-type* cells. In these polarity-impaired cells, signaling mediated by the receptor-type tyrosine phosphatase Ptp10D upon engagement with its ligand Sas in the surrounding *wild-type* cells triggers cell competition via EGFR pathway inhibition and JNK pathway activation, which induces apoptosis of the mutant cells. Here, we investigate whether directly triggering cytoskeletal deregulation (which usually occurs downstream of cell polarity disruptions) is sufficient to trigger their elimination by cell competition via the Sas-Ptp10D signaling system. We show that actin cytoskeleton deregulated cells (as induced by *RhoGEF2* overexpression (*RhoGEF2^OE^*)) are eliminated when surrounded by *wild-type* cells, and that *Ptp10D* knockdown increases *RhoGEF2^OE^* clone growth, revealing the importance of Ptp10D in the elimination of *RhoGEF2^OE^* cells. Mechanistically, in clones that are moderately overexpressing *RhoGEF2^OE^, Ptp10D* knockdown rescued cell elimination by reducing Hippo signaling. In this setting, JNK and EGFR-Ras signaling were not affected (in contrast to what occurs in apico-basal mutant cells), suggesting that Sas-Ptp10D may regulate the Hippo pathway directly in *RhoGEF2^OE^* cells. We also found that mutations in the apical cell polarity protein, Crb, partially rescued the elimination of *RhoGEF2^OE^* clones, showing that Crb normally plays a role in *RhoGEF2^OE^* clone elimination. In this setting, in which *RhoGEF2^OE^* is highly overexpressed, JNK and Hippo signaling were elevated while EGFR-Ras signaling was reduced, and *crb* loss normalized these pathways. *crb* mutant cells also showed reduced abundance of and apical membrane localization of Ptp10D, suggesting that Crb may play an important role in Sas-Ptp10D mediated cell competition. Thus, actin cytoskeleton deregulation, caused by *RhoGEF2^OE^*, results in clone elimination dependent on Crb, Ptp10D, and Hippo signaling. Altogether, our results reveal that Ptp10D is acting more broadly in cell competition to trigger the elimination of actin cytoskeleton deregulated loser cells, as well as polarity-impaired cells.

## Introduction

Cell competition is an intrinsic surveillance mechanism that functions within tissues, where cells gauge the fitness of their neighbors. The “more fit” cells (“winners”) instigate the elimination of “less fit” (“loser”) cells (Morata and Ripoll, 1975; Simpson and Morata, 1981). Importantly, the elimination of these defective (or suboptimal) loser cells is context dependent, such that the loser cells are completely viable when surrounded by cells of the same loser genotype (Morata and Ripoll, 1975; Wodarz, 2000). After cell competition occurs, any space previously occupied by now eliminated loser cells is then filled by winner cells through the process of “compensatory cell proliferation”, maintaining tissue homeostasis (Fan and Bergmann, 2008; Martín et al., 2009). It is now clear that cell competition mechanisms play an important role during development, aging, and cancer (Merino et al., 2016).

The first description of a cell competition mechanism was made in a *Drosophila* model, and involved heterozygous mutations in the ribosomal protein gene, *Minute* (*M*). When *M^+/−^* cell co-existed with other *M^+/−^* cells, no cells were eliminated and the imaginal disc developed normally. However, when *M^+/−^*co-existed with *M^+/+^* cells, *M^+/−^* cells were eliminated from the tissue (Morata and Ripoll, 1975). In this initial cell competition model, it was thought that the mechanism behind cell elimination was differential cell growth rates (Simpson and Morata, 1981). It wasn’t until 20 years later when another key discovery opened the field again, where it was shown that *M^+/−^* clone elimination was not only due to differential growth rates, but was also apoptosis-dependent and regulated by access to the survival factor decapentaplegic/bone morphogenetic protein/tumor growth factor-β (Dpp/BMP/TGF-β) in the wing imaginal disc, demonstrating cell fitness could also depend on the ability of cells to compete for limited survival factors (Abrams, 2002; Moreno et al., 2002).

Interest in cell competition surged further when it was linked to cancer through the discovery of super-competitors, where the mutant cells are winners and *wild-type* cells are losers. The proto-oncogene *dMyc*, a conserved transcription factor that regulates various downstream targets involved in cell growth and ribosome biogenesis (de la Cova and Johnston, 2006; Johnston et al., 1999), was the first gene identified that fit this concept – clones expressing high levels of *dMyc* expand at the expense of the surrounding *wild-type* tissue until they occupied the entire imaginal disc (de la Cova et al., 2004; Moreno and Basler, 2004). This expansion required the elimination of surrounding *wild-type* cells through apoptosis triggered by Dpp deprivation, JNK activation (Moreno and Basler, 2004), and the activation of Hid (the apoptotic inducer) in the loser cells (de la Cova et al., 2004). This discovery was particularly compelling because it resembled the early stages of cancer development (Rhiner and Moreno, 2009). Conversely, downregulation of *dMyc* in clones led to their elimination, mirroring the effects seen with *Minute* mutations (de la Cova et al., 2004; Johnston et al., 1999; Moreno and Basler, 2004).

Another important mechanism that has been shown to induce cell competition are mutations in the core regulators of apico-basal polarity, such as *scribble (scrib)*, *discs large 1* (*dlg1*, hereafter *dlg*), and *lethal (2) giant larvae* (*l(2)gl*, hereafter *lgl*), which produce loser cells that are removed via JNK-dependent apoptosis (Brumby and Richardson, 2003; Chen et al., 2010; Igaki et al., 2009; Menendez et al., 2010; Ohsawa et al., 2011; Tamori et al., 2010). Similarly, clones overexpressing the apical polarity determinant *crumbs* (*crb*) are losers, and are eliminated by apoptosis when surrounded by *wild-type* cells (Hafezi et al., 2012). In contrast, *crb* mutant clones are winners, undergoing increased proliferation and inducing apoptosis in the surrounding *wild-type* cells (Hafezi et al., 2012). *scrib, dlg,* and *lgl* are also recognized as tumor suppressor genes, as mutations in these genes can cause entire tissues to become tumorigenic (Wodarz, 2000) and, therefore, cell competition involving these genes (and other tumor suppressors) is also known as ‘tumor suppressive cell competition’ (Kanda and Igaki, 2020). Other genes in this group include endocytic pathway components, such as *avalanche*/*Syntaxin 7* (*avl*), *Vps25*, *erupted*/*TSG101*, and *Rab5* (Ballesteros-Arias et al., 2014; Menut et al., 2007; Moberg et al., 2005; Thompson et al., 2005; Vaccari and Bilder, 2005).

Hippo signaling is a central pathway for cell competition mechanisms, including those involving *Minute*, *dMyc*, *crb*, and tumor-suppressive cell competition (Chen et al., 2012; Froldi et al., 2010; Hafezi et al., 2012; Menendez et al., 2010; Tyler et al., 2007). This pathway is important in regulating tissue growth and has been implicated in various cancers (Staley and Irvine, 2012; Zhao et al., 2011). The core components of the Hippo pathway include the kinases Hippo and Warts, along with the adaptors Salvador and Mats, which together prevent the nuclear accumulation of the co-transcription factor Yorkie (Yki; known as YAP in mammals). When the Hippo pathway is downregulated, Yki accumulates in the nucleus and binds to the TEAD family transcription factor Scalloped (Sd), leading to the transcription of genes that promote cell proliferation (e.g. the cell cycle gene, *Cyclin E*) and inhibit apoptosis (e.g. *Diap1* (*Drosophila* inhibitor of apoptosis 1)). Mutations in all the core Hippo pathway components result in a super-competitor phenotype in a clonal context (Tyler et al., 2007). Conversely, high levels of Hippo signaling confer a loser cell phenotype. The mechanism by which the Hippo pathway is regulated in different cell competition scenarios has not been precisely discerned. However, in *crb* mutant super-competitor clones, Expanded (an upstream negative regulator of Hippo signaling) is mislocalized, which leads to impaired Hippo signaling (Chen et al., 2010; Robinson et al., 2010). For more information on cell competition mechanisms and signaling pathways see the following reviews (Cong and Cagan, 2024; Fahey-Lozano et al., 2019; Levayer and Moreno, 2013; Nagata and Igaki, 2024).

For many years, a question that remained unanswered in the field of cell competition was how are loser and winner cells communicating with each other to discern the fitness status of neighbouring cells and subsequently orchestrate their elimination? One possible mechanism involves the *Flower* gene, different isoforms of which regulate loser cell status during cell competition (Rhiner et al., 2010). Notably, Flower expression occurs downstream of several fitness modulators, including mutations in *dMyc*, *Minute*, *scrib*, and the Dpp pathway (Merino et al., 2015; Rhiner et al., 2010). Another mechanism is via innate immune signaling pathways, such as Toll-related receptors and NF-κB transcription factors, which are essential for both *Minute* and *dMyc* cell competition via activation of cell death genes to promote loser cell elimination (Alpar et al., 2018; de Beco et al., 2012; Meyer et al., 2014). (Interestingly, the opposite occurs in *scrib* mutants, where Toll activation inhibits cell competition and therefore triggers tumour-like growth (Katsukawa et al., 2018)). A recently discovered mechanism of cell competition involving a direct receptor-ligand interaction is the Sas-Ptp10D axis, which was shown to particular concern loser cells with compromised cell polarity within epithelial tissues (Yamamoto et al., 2017). Ptp10D is a receptor-type tyrosine phosphatase which undergoes re-localization to the lateral membrane in polarity-deficient cells, while its ligand Stranded at second (Sas) is similarly re-localized to the lateral membrane in neighbouring *wild-type* cells. Ptp10D activation via Sas binding inhibits the EGFR-Ras signaling pathway, enabling pro-apoptotic JNK signaling and the elimination of the polarity-deficient cells.

In line with how cells lacking the apico-basal polarity proteins Scrib and Dlg, exhibit a loser phenotype when adjacent to *wild-type* cells (Brumby and Richardson, 2003; Grzeschik et al., 2007; Igaki et al., 2006; Menendez et al., 2010), changes in cell morphology governed by regulators of the actin cytoskeleton also contribute to the elimination of cells by neighboring *wild-type* cells (Brumby et al., 2011; Khoo et al., 2013). Given the intricate regulatory network between polarity proteins and cytoskeletal regulators (Mack and Georgiou, 2014), we sought to investigate whether inducing morphology changes alone by deregulating the actin cytoskeleton, without substantial alterations in cell polarity, could initiate cell competition, and whether this process involved Sas-Ptp10D signaling. Therefore, we examined the involvement of the Sas-Ptp10D cell competition system in *RhoGEF2* overexpressing (*RhoGEF2^OE^*) cells, which leads to actin cytoskeleton deregulation without gross cell polarity effects (Khoo et al., 2013). We show that Sas-Ptp10D signaling also contributes to the elimination of actin cytoskeletal deregulated cells. Mechanistically, in cells with moderate *RhoGEF2* overexpression, the Hippo pathway is activated but the EGFR-Ras and JNK pathways are not affected, and Ptp10D knockdown prevents cell competition by inhibiting the Hippo pathway. Additionally, we show that the apical cell polarity protein, Crb, also plays a role in the elimination of *RhoGEF2^OE^* cytoskeletal deregulated cells via cell competition. In cells with high *RhoGEF2* overexpression, Hippo and JNK signaling were activated, and EGFR-Ras signaling was reduced, and in this scenario *crb* loss normalized these pathways. Finally, we found that *crb* mutant cells exhibit reduced expression and membrane localization of Ptp10D, suggesting Crb may be important for Ptp10D signaling-mediated elimination of *RhoGEF2^OE^* clones. The findings from this study expand on the role of Ptp10D signaling in cell competition and demonstrate that it also plays a role in the elimination of actin cytoskeletal deregulated loser cells.

## Results

### Ptp10D is required for *scrib* mutant clone elimination, but Ptp10D overexpression does not enhance cell competition

The discovery of the role of the Sas-Ptp10D signaling system in polarity-deficient cell competition marked a significant breakthrough (Yamamoto et al., 2017). Prior to this, the mechanisms by which polarity-disrupted cells were recognized and underwent cell competition were not properly understood. However, recent controversy has emerged regarding the obligatory role of Ptp10D in this context. A study conducted by David Bilder’s lab (Gerlach et al., 2022) reported that depleting Ptp10D in *scrib^−/−^* clones, using the same *Ptp10D-RNAi* stock as used in the Yamamoto et al., 2017 study did not rescue clone size in third instar (L3) *Drosophila* eye imaginal discs under their laboratory conditions. Surprised by this finding, Gerlach et al. tested various MARCM clone induction tools, polarity deficiencies, a *Ptp10D* mutant, different food recipes (molasses vs. corn syrup and different protein concentrations), and other husbandry conditions such as crowding (Gerlach et al., 2022). Despite these efforts, they still observed no rescue of clone elimination when co-depleting Ptp10D with an apico-basal polarity regulator, suggesting that Ptp10D might not always be involved in the elimination of polarity-deficient clones. However, another recent study also utilizing the same *Ptp10D-RNAi* stock as used by Yamamoto et al., 2017 found that Ptp10D was necessary for the elimination of polarity disrupted clones (Liu et al., 2022). Consequently, we also attempted to replicate Yamamoto et al.’s results using our own stocks and laboratory conditions.

To evaluate this, we performed a clonal knockdown of *Ptp10D* in polarity-disrupted (*scrib^−/−^*) cells and measured the relative clone size by quantifying the volume of GFP+ tissue relative to the total volume of the disc (see Materials and Methods). We found that *Ptp10D* knockdown (*Ptp10D^KD^*) with *UAS-Ptp10D-RNAi* lines, was able to partially rescue the elimination of *scrib^−/−^*clones [compare Fig 1C and 1D to 1B; quantified in 1E], consistent with the findings of Yamamoto et al., 2017 and Liu et al., 2022. However, we noted that the increase/rescue in clone size was not as pronounced as that reported by the previous studies, and did not reach *wild-type* clone size, a discrepancy which might potentially be explained by the use of different *Ptp10D^KD^* lines in our experiments. However, since no rescue at all was observed in the Gerlach et al., 2022 study, it is possible that other cell competition mechanisms may be at play under their study conditions.

**Figure 1.**
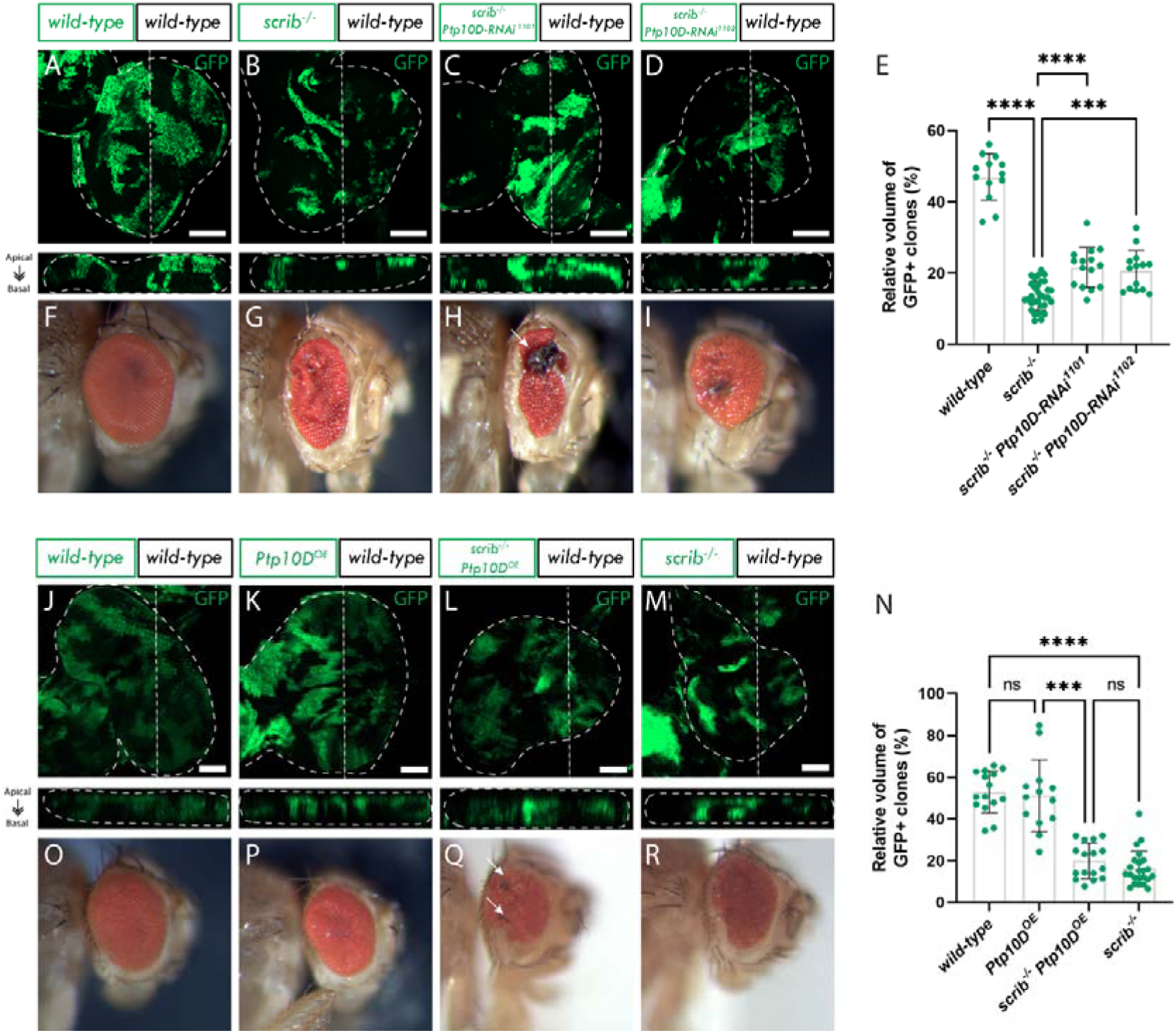
Ptp10D is required for scrib mutant clone elimination and Ptp10D overexpression does not enhance cell competition. A-D, eye discs of ey-FLP-MARCM-induced mosaics (clones are marked by the presence of GFP and wild-type tissue is unmarked): A) wild-type; B) scrib^−/−^; C) scrib^−/−^ UAS-Ptp10D-RNAi^1101^; D) scrib^−/−^ UAS-Ptp10D-RNAi^1102^; E) Quantifications of relative GFP+ clone volume [wild-type (n=13); scrib^−/−^ (n=32); scrib^−/−^ UAS-Ptp10D-RNAi^1101^ (n=15); scrib^−/−^ UAS-Ptp10D-RNAi^1102^ (n=14); Statistical test used were one-way ANOVA, p<0.05, with Tukey’s multiple comparison test. In graph: ****p<0.0001, ***p: 0.0003]; F-I, adult eyes of the corresponding eye imaginal discs (arrow in H indicate necrotic tissue); J-M, eye discs of ey-FLP-MARCM-induced mosaics (clones are marked by the presence of GFP and wild-type tissue is unmarked): J) wild-type; K) Ptp10D overexpression (Ptp10D^OE^); L) scrib^−/−^ Ptp10D^OE^; M) scrib^−/−^. N) Quantifications of relative GFP+ clone volume [wild-type (n=15); Ptp10D^OE^ (n=13); scrib^−/−^ Ptp10D^OE^ (n=16); scrib^−/−^ (n=23); Statistical test used was Kruskal-Wallis test, p<0.05, with Dunn’s multiple comparison test. In graph: ****p<0.0001, ***p: 0.0009]; O-R) adult eyes of the corresponding eye imaginal discs (arrows in Q indicate necrotic speckles). ns = not significant; error bars = SD; scale bars indicate 20 µm. Below each confocal image is an xz cross section of the corresponding eye-antennal disc from the apical (top) to basal (bottom) edge, with the position of the chosen xz sections indicated by a vertical dotted line in the xy images. Dotted lines surrounding discs illustrate disc boundaries.

Notably, we utilized two *Ptp10D^KD^* lines obtained from the Vienna *Drosophila* Research Center (VDRC), numbered 1101 and 1102, different to the studies by Yamamoto et al., 2017, Liu et al. 2022 and Gerlach et al., 2022, which used the *Ptp10D-RNAi* 39086 line from the Bloomington *Drosophila* Stock Center (BDSC). The two *Ptp10D^KD^* lines we used in this study exhibited very similar knockdown of Ptp10D in *scrib^−/−^* third instar larval eye epithelia (around 50%) [compare Supp Figs 1D’ and 1E’ to 1F’; quantified in Supp Fig 1H] (and importantly, the downregulation of *Ptp10D* alone did not affect the total volume of the GFP^+^ tissue when compared to the *wild-type* GFP^+^ tissue in *wild-type* eye-antennal disc mosaics [compare Supp Figs 1B and 1C to 1A; quantified in Supp Fig 1G]). However, we found that the *Ptp10D^KD^* line 1101 induced a stronger adult eye phenotype than the *Ptp10D^KD^* line 1102 in a *scrib* mutant background [compare Figs 1H and 1I]. The expression of *Ptp10D^KD^* 1101 in *scrib^−/−^* clones resulted in significantly smaller adult eyes and a higher percentage of severe adult eye phenotypes, which are characterized by a highly rough eye surface and significant patches of necrotic tissue [indicated by the arrow in Fig 1H]. When the *Ptp10D^KD^*1102 construct was expressed in *scrib^−/−^* clones, the adult eye phenotype was generally less severe, presenting only necrotic speckles instead of large patches of necrotic tissue [Fig 1I]. It is possible that the *Ptp10D^KD^* line 1101 may induce a greater reduction of Ptp10D in *scrib^−/−^* clones at the pupal-adult stages, accounting for the more severe adult eye phenotype. However, for both *Ptp10D^KD^* lines, the *scrib^−/−^ Ptp10D^KD^* adult eyes were similar to that observed by Yamamoto et al., 2017, showing greater disruptions and necrosis than *scrib^−/−^* alone adult eyes, and were contrary to that observed by Gerlach et al., 2022, where no necrotic tissue or further disruption to the *scrib^−/−^* adult eye structure was observed. Since the effect of *Ptp10D^KD^* line 1101 was stronger, it was used for all subsequent experiments.

Furthermore, in support of the Yamamoto et al. study, we found that *Sas^KD^* in surrounding *wild-type* cells also rescued *scrib^−/−^* cell elimination [Supp Fig 2]. Interestingly, Gerlach et al. 2022 conducted the same experiment and although they did observe a similar patchy/necrotic adult eye phenotype as observed by us and Yamamoto et al., 2017, they did not observe an increase in *scrib^−/−^* clone size.

**Figure 2.**
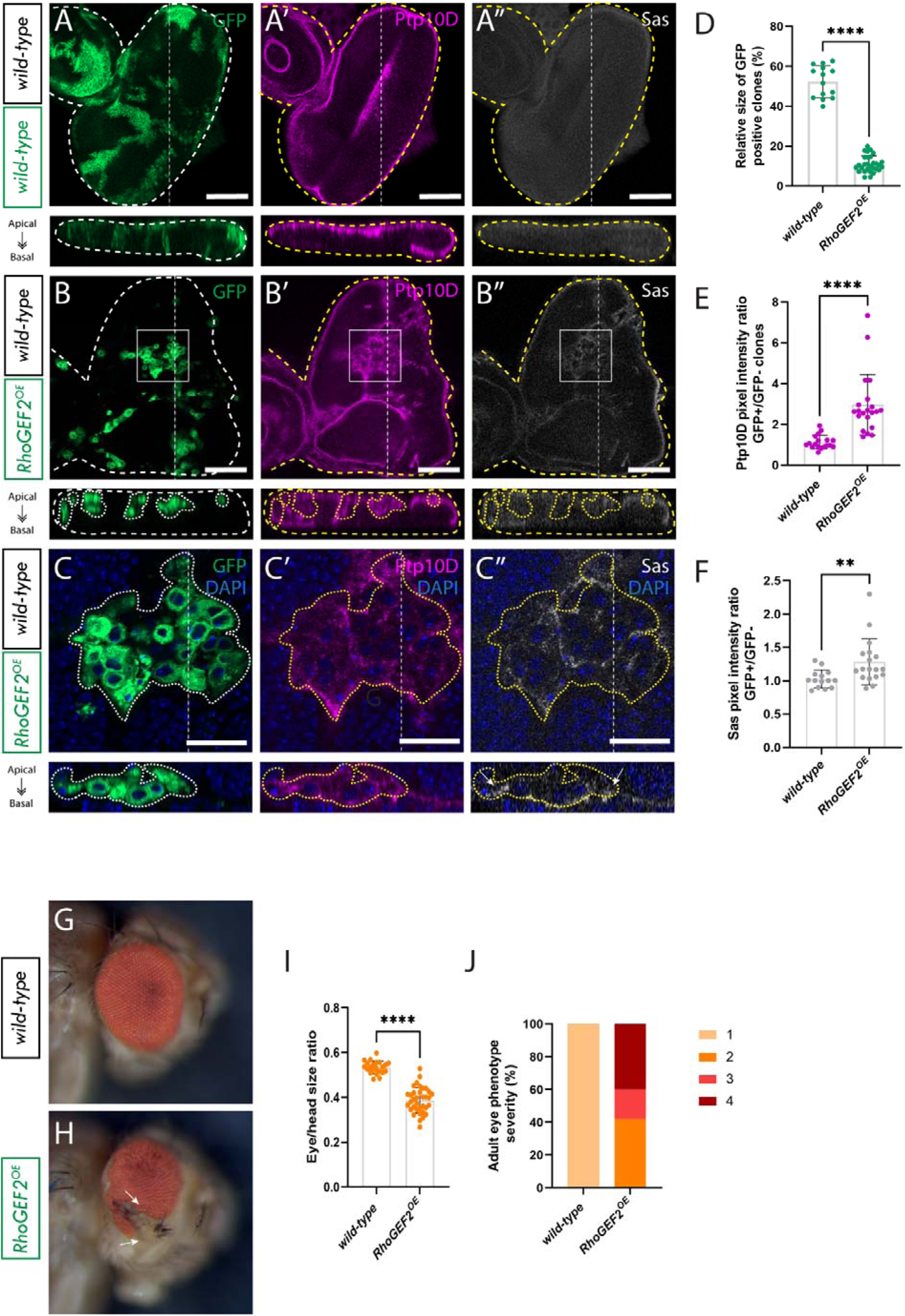
Ptp10D and Sas accumulate at the RhoGEF2^OE^ clone cell membrane, in RhoGEF2^OE^ eye disc mosaics. A) eye disc of wild-type ey-FLP-MARCM-induced mosaics; A’) Ptp10D immunostain; A’’) Sas immunostain; B) Eye disc of RhoGEF2^OE^ ey-FLP-MARCM-induced mosaics (RhoGEF2^OE^ clones are marked by the presence of GFP and wild-type tissue is unmarked); B’) Ptp10D immunostain; B’’) Sas immunostain (rectangles indicate magnified sections shown below); C) Magnified section of eye disc of ey-FLP-MARCM-induced mosaics + DAPI; C’) Ptp10D immunostain + DAPI; C’’) Sas immunostain + DAPI (arrows indicate Sas accumulation); D) Quantification of relative GFP+ clone volume [wild-type (n=14); RhoGEF2^OE^ (n= 29); Statistical test used was unpaired t-test, p<0.05. In graph: ****p<0.0001]; E) Quantification of Ptp10D immunostain pixel intensity ratio between GFP+ and GFP-clones [wild-type (n=14); RhoGEF2^OE^ (n= 18); Statistical test used was Mann-Whitney test, p<0.05. In graph: ****p<0.0001]; F) Quantification of Sas immunostain pixel intensity ratio between GFP+ and GFP-clones [wild-type (n=14); RhoGEF2^OE^ (n= 18); Statistical test used was Mann-Whitney test, p<0.05. In graph: **p: 0.0036]; G) wild-type mosaic adult eye; H) RhoGEF2^OE^ mosaic adult eye (arrows indicate protrusion); I) Relative eye to head size quantification [wild-type (n=22); RhoGEF2^OE^ (n=35), Statistical test used was unpaired t-test, p<0.05. In graph: ****p<0.0001]; J) Percentages of adult eyes with the following phenotypes 1: wild-type, 2: slight rough eye, 3: rougher eye and/or necrotic speckles 4: rougher eye and/or protrusions. ns = not significant; error bars = SD; scale bars in And B indicate 50 µm, scale bars in C indicate 20 µm. Below each confocal image is an xz cross section of the corresponding eye-antennal disc from the apical (top) to basal (bottom) edge, with the position of the chosen xz sections indicated by a vertical dotted line in the xy images. Dotted lines surrounding discs or clones illustrate disc/clone boundaries. Note that folding at the edges of the discs can result in Ptp10D and Sas being observed basally in the xz sections in these regions, despite the staining still being localized at the apical membrane.

Out of curiosity, we tested whether the levels of Ptp10D might be rate-limiting for cell competition of polarity-impaired cells – i.e. does increasing the amount of Ptp10D in *scrib^−/−^*clones induce stronger cell elimination? We observed that overexpression of *Ptp10D* in *scrib^−/−^* clones did not result in any significant further reduction in clone size [compare Figs 1L and 1M; quantified in 1N]. Adult eyes resulting from the *scrib^−/−^* clones overexpressing *Ptp10D* (*Ptp10D^OE^*) are a similar size to *scrib^−/−^* mosaics, but some contain small necrotic speckles [compare Figs 1Q and 1R]. Thus, the levels of Ptp10D in *scrib^−/−^* mutant clones are sufficient to be fully activated by Sas in the adjacent *wild-type* cells.

### Cytoskeletal deregulated clones are eliminated by cell competition

Cytoskeletal deregulation is a process that occurs downstream of cell polarity disruptions (Brumby et al., 2011; Elsum et al., 2012). We therefore sought to investigate if directly deregulating the actin cytoskeleton by overexpressing *RhoGEF2^OE^*, without overtly affecting cell polarity, was sufficient to trigger cell competition. Although our previous studies had revealed that *RhoGEF2^OE^* reduced clone size (Brumby et al., 2011; Khoo et al., 2013), it was not quantified.

As anticipated, expression of *RhoGEF2^OE^* in clones resulted in a significant reduction in the volume of *RhoGEF2^OE^* GFP^+^ tissue, which constituted only an average of 10% of the total eye disc volume, compared to the 50% average volume of *wild-type* GFP^+^ tissue in control eye disc mosaics [compare Figs 2A and 2B; quantified in 2D]. Importantly, it was previously observed that when *RhoGEF2* was overexpressed in the whole eye tissue, these cells did not exhibit reduced growth but instead overgrew (Brumby et al., 2011; Khoo et al., 2013), indicating that *RhoGEF2^OE^* cells are not intrinsically sick, and suggesting it is only in a clonal context that they are eliminated by cell competition. Interestingly *RhoGEF2^OE^* clonal tissue presented with a cyst-like phenotype (Fig 2C). It has previously been described in wing imaginal discs that this rounded ball-like shape is a consequence of recruitment of Actin, Myosin, and Moesin to the interface between cells with different fates, which induces apical constriction and a reduction in lateral contact area between different cells (Bielmeier et al., 2016).

Next, to explore the potential involvement of the Sas-Ptp10D system in the elimination of *RhoGEF2^OE^* clones we assessed the localization of Ptp10D and Sas in *RhoGEF2^OE^* eye disc mosaics by performing Ptp10D and Sas immunostains. The results revealed a notable shift in the localization of Ptp10D and Sas; no longer confined to the apical membrane but instead being found more basally, either within or surrounding the *RhoGEF2^OE^* clones [compare Figs 2A’ and 2A’’ to 2B’ and 2B’’; magnified in C’ and C’’]. Within *RhoGEF2^OE^*clones in eye disc mosaics, quantification showed an approximately 3-fold average increase in Ptp10D abundance [Fig 2E], and just under a 1.5-fold average increase for Sas abundance [Fig 2F]. Furthermore, DAPI staining of cell nuclei and the membrane-bound mCD8-GFP marker highlighted a concentration of Ptp10D and Sas at the cell membranes [Fig 2 C-C’’], further suggesting that Ptp10D and Sas might be playing a role in the elimination of these clones. Moreover, the size of adult eyes resulting from *RhoGEF2^OE^* eye-antennal disc mosaics was significantly diminished compared to *wild-type* adult eyes [compare Figs 2G and 2H; quantified in 2I]. This observation implies that *RhoGEF2^OE^* clones are being eliminated from the tissue at a higher rate than the surrounding *wild-type* cells are able to replace them (through compensatory cell proliferation mechanisms), analogous to the situation observed in *scrib^−/−^* adult eye mosaics. We then also measured the rugosity/deformity of *RhoGEF2^OE^* mosaic adult eye phenotypes on a 4-point scale (where a *wild-type* appearance is 1); 40% exhibited slight tissue rugosity akin to that observed in *scrib^−/−^* adult eye mosaics (rated 2), whilst 20% of *RhoGEF2^OE^* adult eyes displayed more pronounced rugosity and/or necrotic speckles (rated 3), and 40% exhibited epithelial protrusions, some resembling parts of the antennae (rated 4) [compare Figs 2G and 2H; plotted in 2J]. These data show that cytoskeletal disorganization due to *RhoGEF2^OE^*expression markedly impacts eye/antenna structure development. Importantly, our results reveal that *RhoGEF2^OE^* clones undergo cell competition and that Ptp10D and Sas protein localization is altered to be more basal in and surrounding these clones.

### Ptp10D knockdown increases *RhoGEF2^OE^* clone volume

To investigate if Ptp10D is playing a role in *RhoGEF2^OE^* cell competition, we knocked down *Ptp10D* in *RhoGEF2^OE^* clones and measured the volume of the *RhoGEF2^OE^* GFP^+^ tissue. If Ptp10D is indeed a critical mediator of *RhoGEF2^OE^* clone elimination, then the downregulation of *Ptp10D* would be expected to rescue the elimination of *RhoGEF2^OE^*clones. To account for *GAL4/UAS* dosage effects and ensure equivalent expression levels of *RhoGEF2^OE^*, a control *UAS* construct (*UAS-Dicer2*) was introduced into the *UAS-RhoGEF2^OE^* stock. Before analyzing clone sizes, it was crucial to confirm Ptp10D upregulation in *RhoGEF2^OE^ Dicer2* clones, since the levels of *RhoGEF2^OE^* in this case are reduced to half, which we considered to be ‘moderate overexpression’. As suspected, the upregulation of Ptp10D was milder than with ‘high overexpression’ of *RhoGEF2* [Fig 2], but a significant 1.5-fold increase in Ptp10D abundance was still observed [Supp Fig 3]. Concurrently, effective Ptp10D knockdown was confirmed when *Ptp10D^KD^* was expressed alone (with a *UAS-myr-RFP* dosage control) and in combination with *RhoGEF2^OE^*, resulting in an average 50% reduction in Ptp10D antibody staining intensity [Supp Fig 3]. Quantification of GFP^+^ tissue volumes revealed that *RhoGEF2^OE^ Dicer2* clones did not show significant size reduction compared to *wild-type* eye disc mosaics [compare Figs 3A and 3B; quantified in 3I], in contrast to the observations with high *RhoGEF2^OE^* expression (no *Dicer2*) [Fig 2]. However, *Ptp10D^KD^ RhoGEF2^OE^* clones were significantly increased in volume compared with *RhoGEF2^OE^ Dicer2* clones [compare Figs 3B and 3C; quantified in 3I], indicating that Ptp10D is playing a role in controlling *RhoGEF2^OE^*clone size. Importantly, Ptp10D knockdown (again together with *UAS-myr-RFP*, for dosage compensation) did not increase the volume of *wild-type* cells [compare Figs 3A and 3D; quantified in 3I]. Taken together, these results indicate that Ptp10D has no effect in *wild-type* cells, but functions in *RhoGEF2^OE^* clones to limit their growth/reduce their competitiveness. Following on from the findings of the Bilder lab (Gerlach et al., 2022) we decided to perform this experiment using a different food recipe. We utilized a low protein food recipe (instead of our regular high yeast, molasses-based food recipe), since we hypothesized that it might make *wild-type* cells more competitive relative to cytoskeletal deregulated cells. Interestingly, when we performed this experiment in low protein food, *RhoGEF2^OE^ Dicer2* clones were significantly smaller than the *wild-type* controls [Supp Fig 4], reiterating that environmental conditions are key to cell competition processes (Agrawal et al., 2016; Gerlach et al., 2022; Sanaki et al., 2020). Furthermore, *Ptp10D* knockdown in *RhoGEF2^OE^* clones, in this setting, rescued the reduced clone size [Supp Fig 4], indicating that Ptp10D is involved in cell competition of *RhoGEF2^OE^*clones. However, since we were interested in the clonal size increase of the *RhoGEF2^OE^ Ptp10D^KD^* clones compared to *RhoGEF2^OE^ Dicer2*, we continued to utilize our regular high yeast, molasses-based food for subsequent experiments.

**Figure 3.**
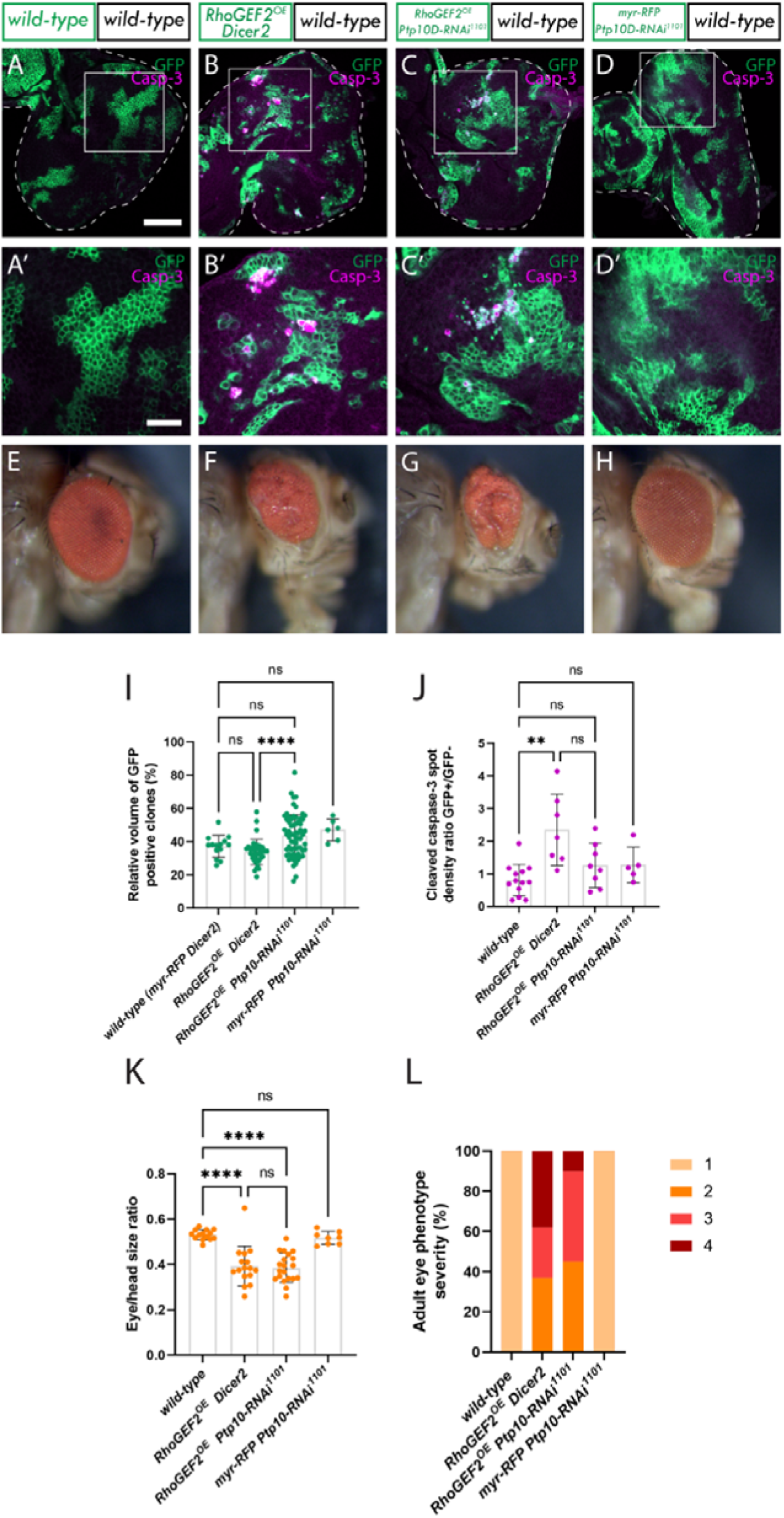
Ptp10D knockdown increases RhoGEF2^OE^ clonal growth, in RhoGEF2^OE^ eye disc mosaics. A-D, eye discs of ey-FLP-MARCM-induced mosaics (clones are marked by the presence of GFP and wild-type tissue is unmarked): A) wild-type; B) UAS-RhoGEF2^OE^ UAS-Dicer-2; C) UAS-RhoGEF2 UAS-Ptp10D-RNAi (VDRC 1102 line); D) UAS-myr-RFP UAS-Ptp10D-RNAi (VDRC 1101 line), showing cleaved caspase-3 immunostains, which is an indicator of cell death by apoptosis (white squares indicate magnified sections shown below). A’-D’) Magnified sections of eye discs of ey-FLP-MARCM-induced mosaics showing cleaved caspase-3 immunostains. E-H) Adult eyes of the corresponding eye imaginal discs; I) Quantification of the relative GFP+ clone volume [wild-type (n=14); RhoGEF2^OE^ Dicer2 (n=36); RhoGEF2 Ptp10D-RNAi 1101 (n=65); myr-RFP UAS-Ptp10D-RNAi 1101 (n=5); Statistical test used were one-way ANOVA, p<0.05, with Tukey’s multiple comparison test. In graph: ****p<0.0001]; J) Cleaved caspase-3 spot density ratio, which normalizes the amount of cleaved caspase-3 positive cells to the total amount of GFP tissue [wild-type (n=13); RhoGEF2^OE^ Dicer2 (n=7); RhoGEF2 Ptp10D-RNAi 1101 (n=8); myr-RFP UAS-Ptp10D-RNAi 1101 (n=5); Statistical test used were Kruskal-Wallis test, p<0.05, with Dunn’s multiple comparison test. In graph: **p:0.0028]; K) Relative eye to head size quantification [wild-type (n=14); RhoGEF2^OE^ Dicer2 (n=16); RhoGEF2 Ptp10D-RNAi 1101 (n=19); myr-RFP UAS-Ptp10D-RNAi 1101 (n=8); Statistical test used were Kruskal-Wallis test, p<0.05, with Dunn’s multiple comparison test. In graph: ****p<0.0001]; L) Percentages of adult eyes for each genotype that show 1: normal phenotype, 2: slight rough eye, 3: rougher eye and/or necrotic speckles, 4: rougher eye and/or protrusions. ns = not significant; error bars = SD; scale bars in A to D indicate 50 µm; scale bars in A’ to D’ indicate 20 µm. Dotted lines surrounding discs illustrate disc boundaries.

**Figure 4.**
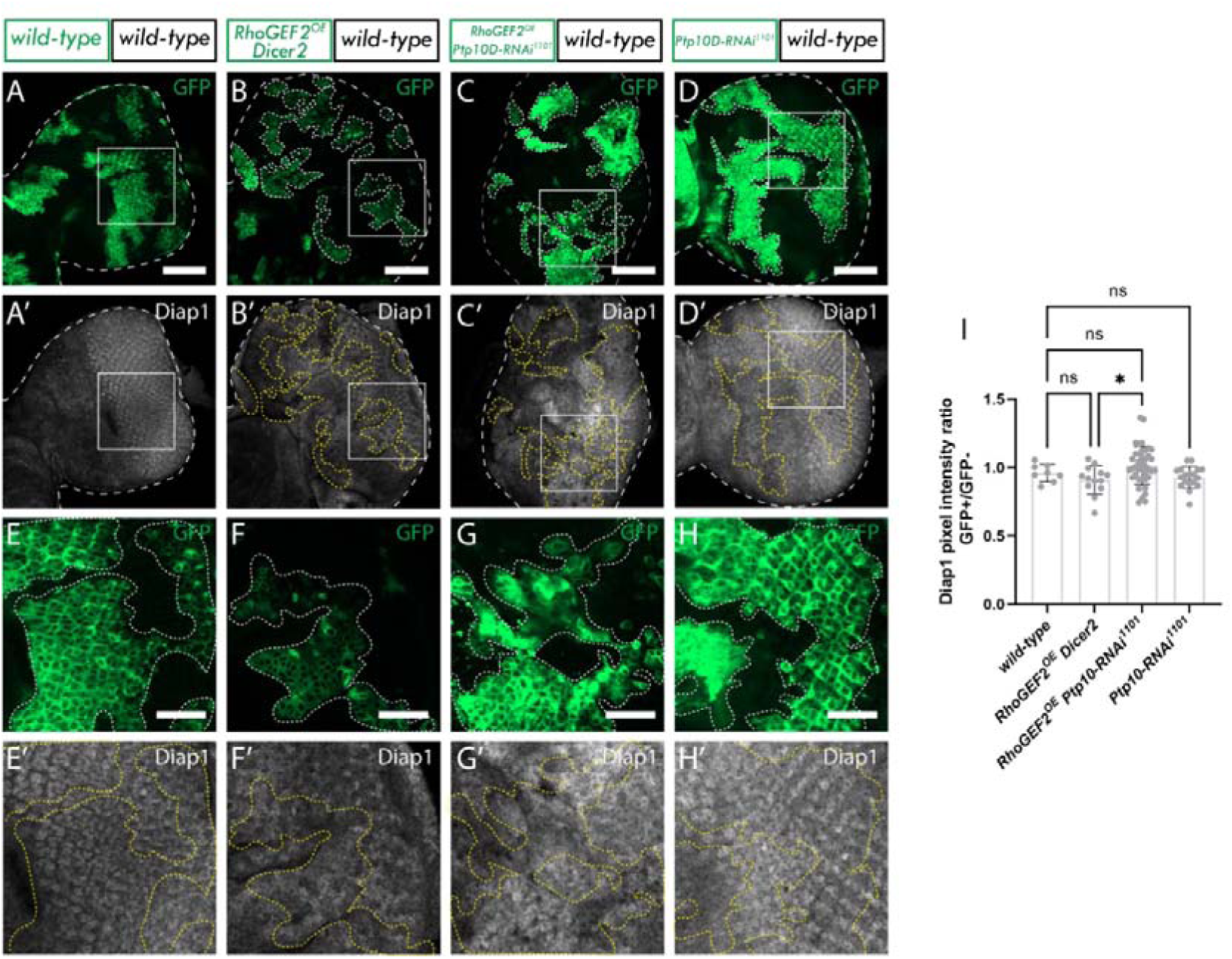
Ptp10D knockdown reduces Hippo signalling in RhoGEF2^OE^ clones. A-D, eye discs of ey-FLP-MARCM-induced mosaics (clones are marked by the presence of GFP and wild-type tissue is unmarked): A) wild-type; B) UAS-RhoGEF2^OE^ UAS-Dicer-2; C) UAS-RhoGEF2 UAS-Ptp10D-RNAi (VDRC 1102 line); D) UAS-myr-RFP UAS-Ptp10D-RNAi (VDRC 1101 line); A’-D’) Diap1 immunostains from A-D; white squares indicate magnified sections shown below. E-H) Magnified sections of eye discs from A-D. E’-F’) Diap1 immunostains from E-F; I) Quantification of Diap1 pixel intensity ratio between GFP+ and GFP-clones [wild-type (n=8); RhoGEF2^OE^ Dicer2 (n=13); RhoGEF2 Ptp10D-RNAi 1101 (n=40); myr-RFP UAS-Ptp10D-RNAi 1101 (n=18); Statistical test used were one-way ANOVA, p<0.05, with Tukey’s multiple comparison test. In graph: *p:0.0323]. ns = not significant; error bars = SD; scale bars in A to D indicate 50 µm; scale bars in E to H indicate 20 µm. Dotted lines surrounding discs or clones illustrate disc/clone boundaries.

Next, we explored cell death in *RhoGEF2^OE^ Dicer2* and *RhoGEF2^OE^ Ptp10D^KD^* clones. Initially, we conducted an immunostain for the active (cleaved) form of the apoptotic protein Caspase-3, a marker for cells undergoing apoptosis. We measured the spot density ratio of cleaved Caspase-3-positive cells between GFP+/GFP− cells, which normalizes the number of positively marked cells to the tissue volume. A significant increase in cleaved Caspase-3 spot density was observed in *RhoGEF2^OE^ Dicer2* clones compared to *wild-type* cells [compare Figs 3A, A’ and Figs B, B’; quantified in 3J]. This result was somewhat unexpected, as no significant decrease in the total volume of *RhoGEF2^OE^ Dicer2* clones (in our regular high yeast, molasses-based food) was observed compared to the *wild-type*. This discrepancy suggests that although these clones undergo cell competition-induced cell death, compensatory cell proliferation mechanisms might be at play in *RhoGEF2^OE^ Dicer2* tissue to offset any clonal size reduction. When *Ptp10D* was knocked down in *RhoGEF2^OE^* clones, no significant decrease in cleaved Caspase-3 spot density ratio was observed compared to *RhoGEF2^OE^ Dicer2* clones, but a downward trend was noted [compare Figs 3B, B’ and Figs 3C, C’; quantified in 3J]. However, importantly, cleaved Caspase-3 levels in *RhoGEF2^OE^ Ptp10D^KD^*clones compared to the *wild-type* clones were not statistically different [compare Figs 3A, A’ and Figs C, C’; quantified in 3J], suggesting that *RhoGEF2^OE^ Ptp10D^KD^* clones undergo less cell death than *RhoGEF2^OE^*. Additionally, *Ptp10D^KD^* clones showed no differences in cleaved Caspase-3 spot density compared to *wild-type*, indicating that Ptp10D knockdown alone does not affect cell death [compare Figs 3A, A’ and Figs D, D’; quantified in 3J].

The size of adult eyes was significantly reduced in both *RhoGEF2^OE^ Dicer2* [Fig 3F] and *RhoGEF2^OE^ Ptp10D^KD^* [Fig 3G], with no significant differences between them, though each were significantly different to the *wild-type* [Fig 3E; quantified in 3K]. However, a higher percentage of *RhoGEF2^OE^ Dicer2* adult eyes exhibited deeper folds and more severe rugosities than *RhoGEF2^OE^ Ptp10D^KD^* adult eyes, suggesting that *Ptp10D* knockdown partially rescues the severe eye phenotype of *RhoGEF2^OE^* mosaics - the opposite of what occurs in the polarity-deficient *Ptp10D^KD^* scenario, where *Ptp10D^KD^* increases the severity of the adult eye phenotype [compare Figs 1H and 3G]. In summary, these findings indicate that Ptp10D plays a role in limiting the growth of (or reducing the competitiveness of) *RhoGEF2^OE^* clones, as *Ptp10D^KD^*leads to increased *RhoGEF2^OE^* clone size and partially rescues the adult eye phenotype.

### *Ptp10D* knockdown reduces Hippo signaling in *RhoGEF2^OE^* clones

The Hippo pathway serves to negatively regulate tissue growth by inhibiting cell proliferation and promoting cell death and, when downregulated, fosters cell proliferation and inhibits cell death, leading to increased tissue size (Tapon et al., 2002). Given that *RhoGEF2^OE^ Ptp10D^KD^* clones exhibited increased size, it was anticipated that these clones would have low levels of Hippo pathway activation. To assess the impact of Ptp10D knockdown on the Hippo pathway in *RhoGEF2^OE^* clones, we stained for the Hippo pathway target Diap1 (Yoo et al., 2002). As expected, Diap1 was found to be upregulated in *RhoGEF2^OE^ Ptp10D^KD^* clones compared to *RhoGEF2^OE^ Dicer2* clones [compare Figs 4B’’ and F’G’ to 4C’ and 4G’; quantified in 4I], indicating impaired Hippo pathway signaling. No differences in Diap1 levels were observed between *RhoGEF2^OE^ Dicer2* and the *wild-type* control [Fig 4A’, E’] or *Ptp10D-RNAi* alone [Fig 4D’, H’].

In the canonical model, Hippo pathway inactivation in *scrib^−/−^ Ptp10D^KD^* clones is a consequence of an F-actin accumulation, which depends on the activation of JNK and EGFR-Ras signaling pathways (Yamamoto et al., 2017). Therefore, to further understand how Hippo is being inactivated in our model we investigated these pathways. Interestingly, we found that while F-actin accumulated in moderate *RhoGEF2^OE^* clones it was not significantly altered upon *Ptp10D* knockdown in moderate *RhoGEF2^OE^* clones [Supp Fig 5], indicating that Hippo inactivation is not triggered by the hyper-accumulation of F-actin. Next, we investigated the JNK signaling pathway - which plays a pivotal role in many types of cell competition (La Marca and Richardson, 2020; Pinal et al., 2019) and is upregulated when *RhoGEF2* is highly overexpressed (without *UAS* dosage control) (Khoo et al., 2013). However, in moderate *RhoGEF2^OE^* clones (with *Dicer2*), we observed no significant increase of the JNK signaling pathway target Mmp1 [Supp Fig 6]. This observation might be attributed to lower levels of *RhoGEF2^OE^* due to the titration of GAL4 transcription factor activity upon introducing a dosage control *UAS* transgene and, therefore, *RhoGEF2* expression levels might be insufficient to trigger JNK pathway activation and concomitant cell elimination. Since no JNK pathway upregulation was observed in this setting, we did not investigate whether *Ptp10D*^KD^*RhoGEF2^OE^* clones affected this signaling pathway, concluding that JNK pathway modulation was unlikely to be a core driver of the increased size of *RhoGEF2^OE^ Ptp10D^KD^*clones. Furthermore, via immunostaining, we did not observe any significant changes in the levels of the phosphorylated (active) form of ERK (pERK; a.k.a. rolled (rl) or MAPK), the canonical downstream effector of Ras signaling. In fact, pERK levels, in *RhoGEF2^OE^ Ptp10D^KD^*, clones were significantly slightly decreased compared to the *wild-type* control [Supp Fig 7], showing that Ptp10D does not inhibit EGFR-Ras signaling in this context. Altogether, these results suggest that the Hippo pathway is being inactivated via an unknown mechanism in *RhoGEF2^OE^ Ptp10D^KD^* clones, where JNK signaling is not hyperactive, EGFR-Ras signaling is not activated, and F-actin hyperaccumulation is not present.

**Figure 5.**
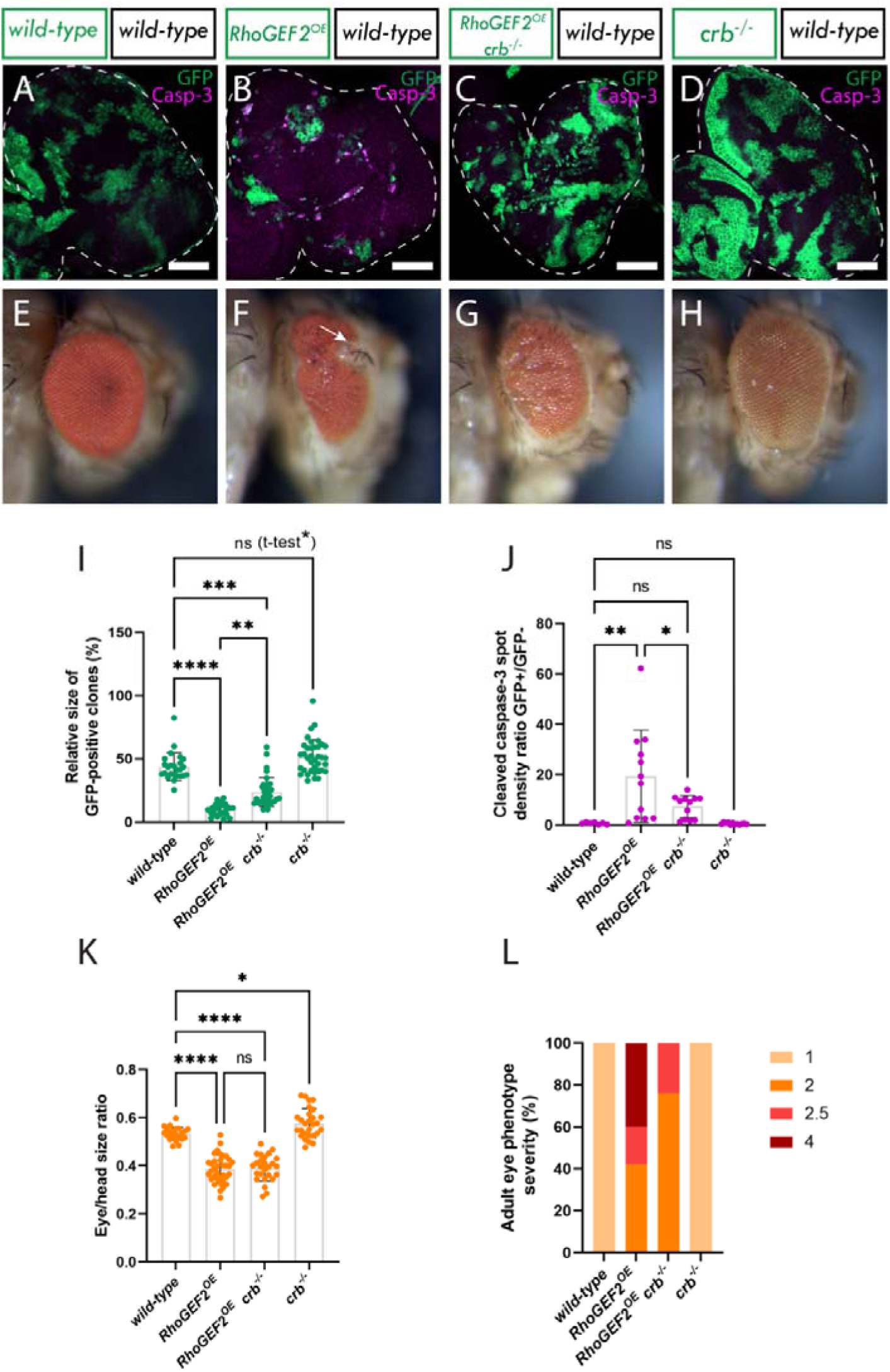
crb mutation results in a partial rescue of RhoGEF2^OE^ clone elimination in eye disc mosaics. A-D, eye discs of ey-FLP-MARCM-induced mosaics (clones are marked by the presence of GFP and wild-type tissue is unmarked): A) wild-type; B) RhoGEF2^OE^; C) UAS-RhoGEF2^OE^ crb^−/−^; D) crb^−/−^, showing cleaved Caspase-3 immunostains, which is an indicator of cell death by apoptosis. E-H) Adult eyes of the corresponding eye imaginal discs. I) Quantification of the relative GFP+ clone volume [wild-type (n=25); RhoGEF2^OE^ (n=25); RhoGEF2 crb^−/−^ (n=34); crb^−/−^ (n=37); Statistical test used were Kruskal-Wallis test, p<0.05, with Dunn’s multiple comparison test. In graph: **p: 0.081; ***p:0.0009; ****p<0.0001. [Since we wanted to look if crb^−/−^ clone increased in volume compared only to the wild-type we performed and unpaired t-test between wild-type and crb^−/−^; p*: 0.0147]; J) Cleaved caspase-3 spot density ratio, which normalizes the amount of cleaved Caspase-3 positive cells to the total amount of GFP tissue [wild-type (n=7); RhoGEF2^OE^ (n=12); RhoGEF2 crb^−/−^ (n=12); crb^−/−^ (n=12); Statistical test used were one-way ANOVA, p<0.05, with Tukey’s multiple comparison test. In graph: *p: 0.0261; **p:0.0019]; K) Relative eye to head size quantification [wild-type (n=22); RhoGEF2^OE^ (n=35); RhoGEF2 crb^−/−^ (n=26); crb^−/−^ (n=28); Statistical test used were one-way ANOVA, p<0.05, with Tukey’s multiple comparison test. In graph: *p:0.0269; ****p<0.0001]; L) Percentages of adult eyes for each genotype that show 1: normal phenotype, 2: slight rough eye, 3: rougher eye and/or necrotic speckles, 4: rougher eye and/or protrusions. ns = not significant; error bars = SD; scale bars indicate 50 µm. Dotted lines surrounding discs illustrate disc boundaries.

**Figure 6.**
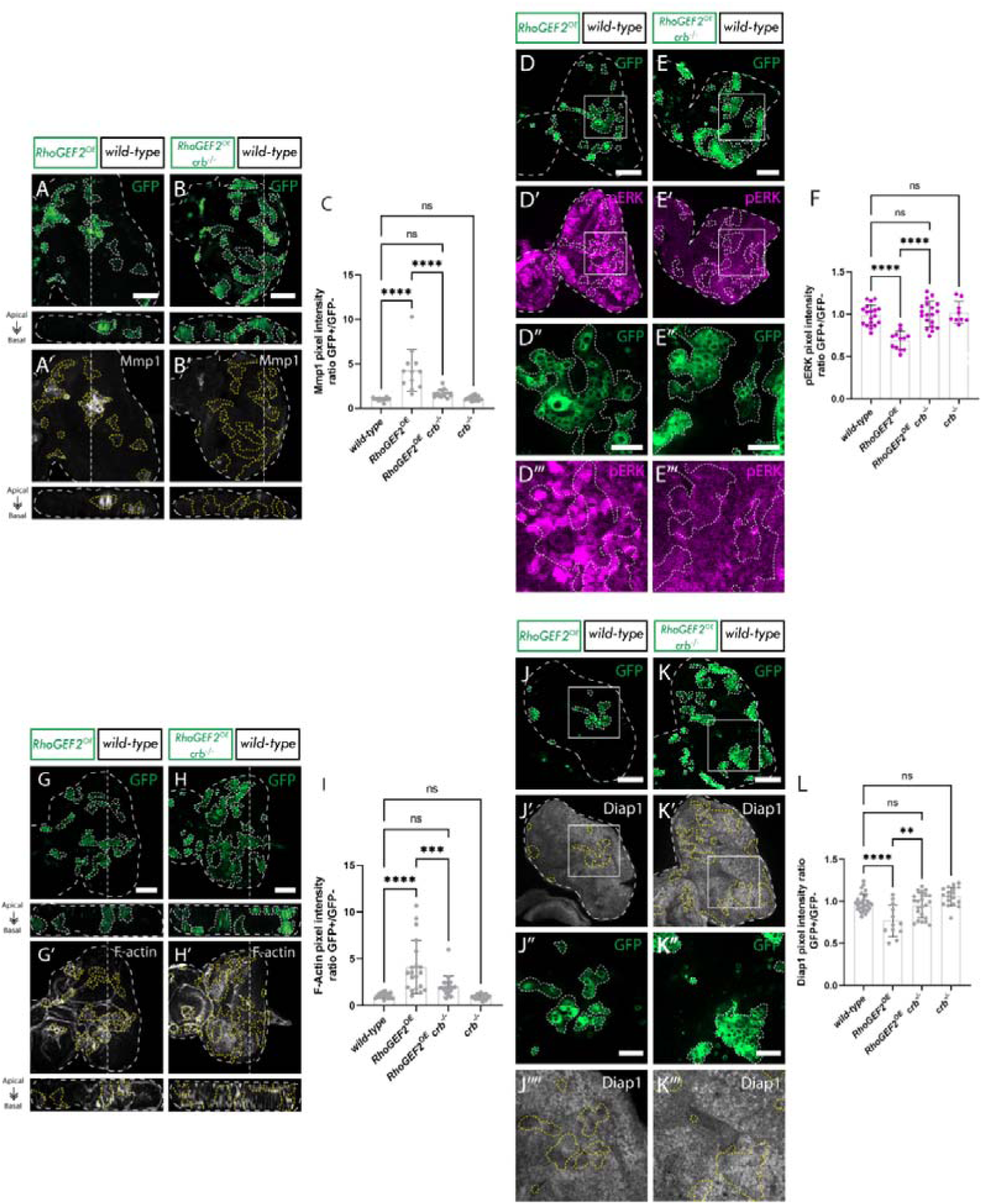
crb loss decreases apical F-Actin accumulation, rescues the elevated JNK signalling, rescues Ras signalling downregulation and reduces the elevated Hippo signalling in RhoGEF2^OE^ clones. A-B, Eye discs of ey-FLP-MARCM-induced mosaics (clones are marked by the presence of GFP and wild-type tissue is unmarked): A) RhoGEF2^OE^; B) RhoGEF2^OE^ crb^−/−^; A’-B’) Mmp1 immunostains from A-B; C) Quantification of Mmp1 pixel intensity ratio between GFP+ and GFP-clones [wild-type (n=18); RhoGEF2^OE^ (n=12); RhoGEF2 crb^−/−^ (n=15); crb^−/−^ (n=18); Statistical test used were one-way ANOVA, p<0.05, with Tukey’s multiple comparison test. In graph: ****p<0.0001]; D-E, Eye discs of ey-FLP-MARCM-induced mosaics (clones are marked by the presence of GFP and wild-type tissue is unmarked): D) RhoGEF2^OE^; E) RhoGEF2^OE^ crb^−/−^; D’-E’) pERK immunostains from D-E; white squares indicate magnified sections shown in panels below; D’’-E’’) Show magnified panels of D-E; D’’’-E’’’) Show pERK immunostains from D’’-E’’. F) Quantification of pERK pixel intensity ratio between GFP+ and GFP− clones [wild-type (n=18); RhoGEF2^OE^ (n=12); RhoGEF2 crb^−/−^ (n=19); crb^−/−^ (n=9); Statistical test used were one-way ANOVA, p<0.05, with Tukey’s multiple comparison test. In graph: ****p<0.0001]; G-H, Eye discs of ey-FLP-MARCM-induced mosaics (clones are marked by the presence of GFP and wild-type tissue is unmarked): G) RhoGEF2^OE^; H) RhoGEF2^OE^ crb^−/−^; G’-H’) F-actin stains; I) Quantification of F-Actin pixel intensity ratio between GFP+ and GFP− clones [wild-type (n=18); RhoGEF2^OE^ (n=20); RhoGEF2 crb^−/−^ (n=18); crb^−/−^ (n=18); Statistical test used were one-way ANOVA, p<0.05, with Tukey’s multiple comparison test. In graph: ***p<0.0008; ****p<0.0001]; J-K) Eye discs of ey-FLP-MARCM-induced mosaics (clones are marked by the presence of GFP and wild-type tissue is unmarked): J) RhoGEF2^OE^; K) RhoGEF2^OE^ crb^−/−^; J’-K’) Diap1 immunostains from J-K; white squares indicate magnified sections shown in panels below; J’’-K’’) Show magnified panels of J-K; J’’’-K’’’) Show Diap1 stains from J’’-K’’. L) Quantification of pERK pixel intensity ratio between GFP+ and GFP− clones [wild-type (n=28); RhoGEF2^OE^ (n=12); RhoGEF2 crb^−/−^ (n=23); crb^−/−^ (n=18); Statistical test used were one-way ANOVA, p<0.05, with Tukey’s multiple comparison test. In graph: **: 0.0013; ****p<0.0001]. Controls have been left out of images for simplicity but can be found in Supp Fig 8. Dotted lines surrounding discs or clones illustrate disc/clone boundaries. ns = not significant; error bars = SD; A, B, D, E, G, H, J, K scale bars indicate 50 µm. D’’, E’’, J’’, K’’ scale bars indicate 20 µm. In A, A’, B, B’, G, G’, H and H’, images below show an xz cross section of the corresponding eye-antennal disc from the apical (top) to basal (bottom) edge, with the position of the chosen xz sections indicated by a vertical dotted line in the xy images.

**Figure 7.**
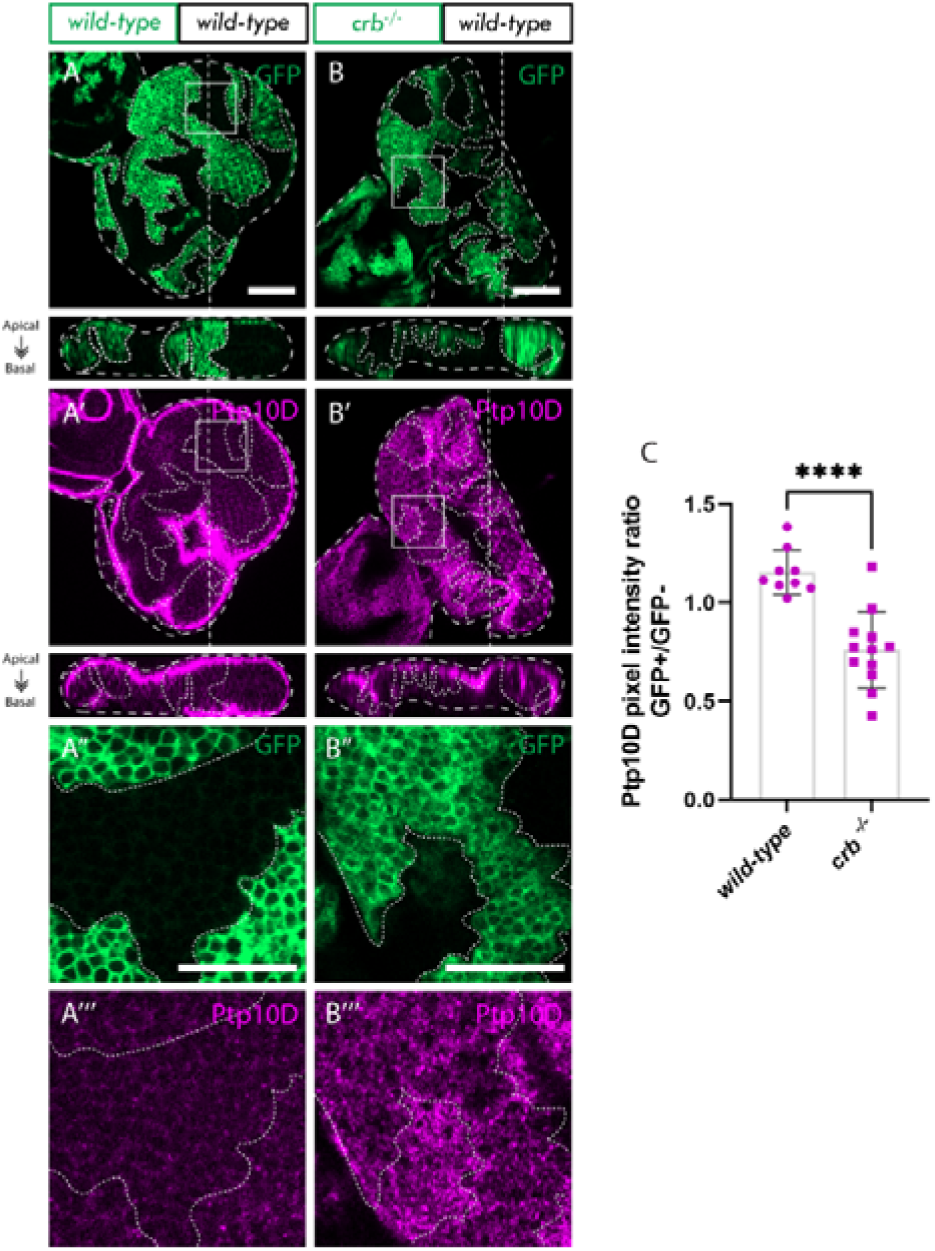
Ptp10D is downregulated in crb mutant clones. A-B, eye discs of ey-FLP-MARCM-induced mosaics (clones are marked by the presence of GFP and wild-type tissue is unmarked): A) wild-type; B) crb^−/−^; A’-B’) Ptp10D immunostains from A-B; white squares indicate magnified sections shown in panels below; A’’-B’’) Magnified sections from A-B; A’’’-B’’’) show Ptp10D immunostains from A’’-B’’. C) Quantification of Ptp10D immunostain pixel intensity ratio between GFP+ and GFP− clones [wild-type (n=9); crb^−/−^ (n= 12); Statistical test used was unpaired t-test, p<0.05. In graph: ****p<0.0001]. ns = not significant; error bars = SD; A, A’, B, B’, scale bars indicate 50 µm. A’’, A’’’, B’’ and B’’’ scale bars indicate 20 µm. Images below show an xz cross section of the corresponding eye-antennal disc from the apical (top) to basal (bottom) edge, with the position of the chosen xz sections indicated by a vertical dotted line in the xy images. Note that folding at the edges of the discs can result in Ptp10D being observed basally in the xz sections in these regions, despite the staining still being localized at the apical membrane. Dotted lines surrounding discs or clones illustrate disc/clone boundaries.

### *crb* mutation partially rescues *RhoGEF2^OE^* clone elimination in eye disc mosaics

Since the apical cell polarity protein Crumbs (Crb) is involved in Hippo pathway regulation and in cell competition (Hafezi et al., 2012; Robinson et al., 2010), we wanted to determine whether *crb* was playing a role in cell competition in *RhoGEF2^OE^* clones. As shown in Fig 2, high *RhoGEF2^OE^*clones are reduced in GFP+ tissue volume (∼10%) compared to *wild-type* clones (∼50%). However, when *crb* was depleted (using the null allele *crb^11A22^*) in *RhoGEF2^OE^* clones, the average GFP+ tissue volume increased to ∼23% [compare Figs 5B and 5C; quantified in 5I], indicating a partial rescue of the *RhoGEF2^OE^* clone elimination phenotype. Additionally, comparing *crb^−/−^* clones and *wild-type* clones revealed a slight but significant increase in volume [compare Figs 5A and 5D; quantified in 5I], consistent with previous reports (Hafezi et al., 2012). The sizes of adult eyes resulting from *RhoGEF2^OE^* and *RhoGEF2^OE^ crb^−/−^* mosaics were not significantly different to one another but were each significantly decreased compared to *wild-type* adult eyes, [compare Figs 5E, 5F and 5G; quantified in 5K], showing that *crb* loss does not rescue the adult eye size decrease in *RhoGEF2^OE^* mosaics. Furthermore, *crb^−/−^* mosaic adult eyes were also significantly larger than *wild-type* adult eyes, consistent with the observation that *crb^−/−^* clones overgrow [compare Figs 5A and 5H; quantified in 5K].

*RhoGEF2^OE^* mosaic adult eyes displayed epithelial protrusions in ∼40% of cases [Fig 5F; quantified in 5L], possibly due to the deregulated actin cytoskeleton. Intriguingly, similar to observations from *RhoGEF2^OE^ Ptp10D*^KD^ clones, the severity of the adult eye phenotypes in *RhoGEF2^OE^ crb^−/−^* mosaics was reduced, with 80% showing only mild rugosities (rated 2), and no protrusions observed in the remaining 20% [Fig 5G; quantified in 5L]. Taken together, these data suggest that Crb is necessary for the elimination of *RhoGEF2^OE^* clones and that although *crb* loss in *RhoGEF2^OE^* clones did not impact adult eye size, it modulated the severity of the *RhoGEF2^OE^*adult eye phenotype.

To investigate whether the loss of *crb* reduced cell death in *RhoGEF2^OE^* clones, we conducted immunostaining for cleaved Caspase-3. Spot density ratio of cleaved caspase-3-positive cells between GFP+ and GFP− cells was measured and, as anticipated, we observed a significantly higher spot density ratio in GFP+ clones in *RhoGEF2^OE^* clones alone compared to *wild-type* [compare Figs 5A and 5B; quantified in 5J], indicating increased cell death. Conversely, *crb^−/−^ RhoGEF2^OE^* clones exhibited significantly fewer cleaved Caspase-3-positive cells, down to *wild-type* levels [compare Figs 5B and 5C; quantified in 5J], indicating that Crb is involved in cell death of *RhoGEF2^OE^* clones through cell competition. Moreover, *crb^−/−^* clones alone had a similar number of cleaved Caspase-3-positive cells to the *wild-type* control [compare Figs 5A and 5D; quantified in 5J], consistent with their resistance to elimination by cell competition.

### *crb* loss decreases apical F-actin accumulation, partially rescues the elevated JNK signaling, rescues EGFR-Ras signaling downregulation, and reduces the elevated Hippo signaling in *RhoGEF2^OE^* clones

To investigate whether the loss of *crb* in *RhoGEF2^OE^*clones is inducing cell death by inhibiting the JNK signaling pathway, we performed immunostaining for the JNK pathway target Mmp1. As expected, we found that Mmp1 was elevated in *RhoGEF2^OE^* clones relative to *wild-type* control clones [Fig 6A’ (and control in Supp Fig 8)], and upon *crb* loss in *RhoGEF2^OE^* clones, Mmp1 levels showed a significant reduction compared to *RhoGEF2^OE^* alone, towards *wild-type* levels [compare Figs 6A’ and 6B’; quantified in 6C], suggesting that Crb contributes to the upregulation of JNK signaling in *RhoGEF2^OE^* clones. Note that in this case we utilized high *RhoGEF2^OE^* and found Mmp1 levels were increased, consistent with previous work (Khoo et al., 2013), which is in contrast to what we observed with only moderate *RhoGEF2^OE^*, where no significant effect on Mmp1 expression was observed [Fig 4]. Next, we investigated whether other pathways involved in the canonical Sas-Ptp10D signaling cascade were affected. Therefore, we assessed the involvement of the EGFR-Ras signaling pathway in the growth of *RhoGEF2^OE^ crb^−/−^* clones by immunostaining for pERK.

**Figure 8.**
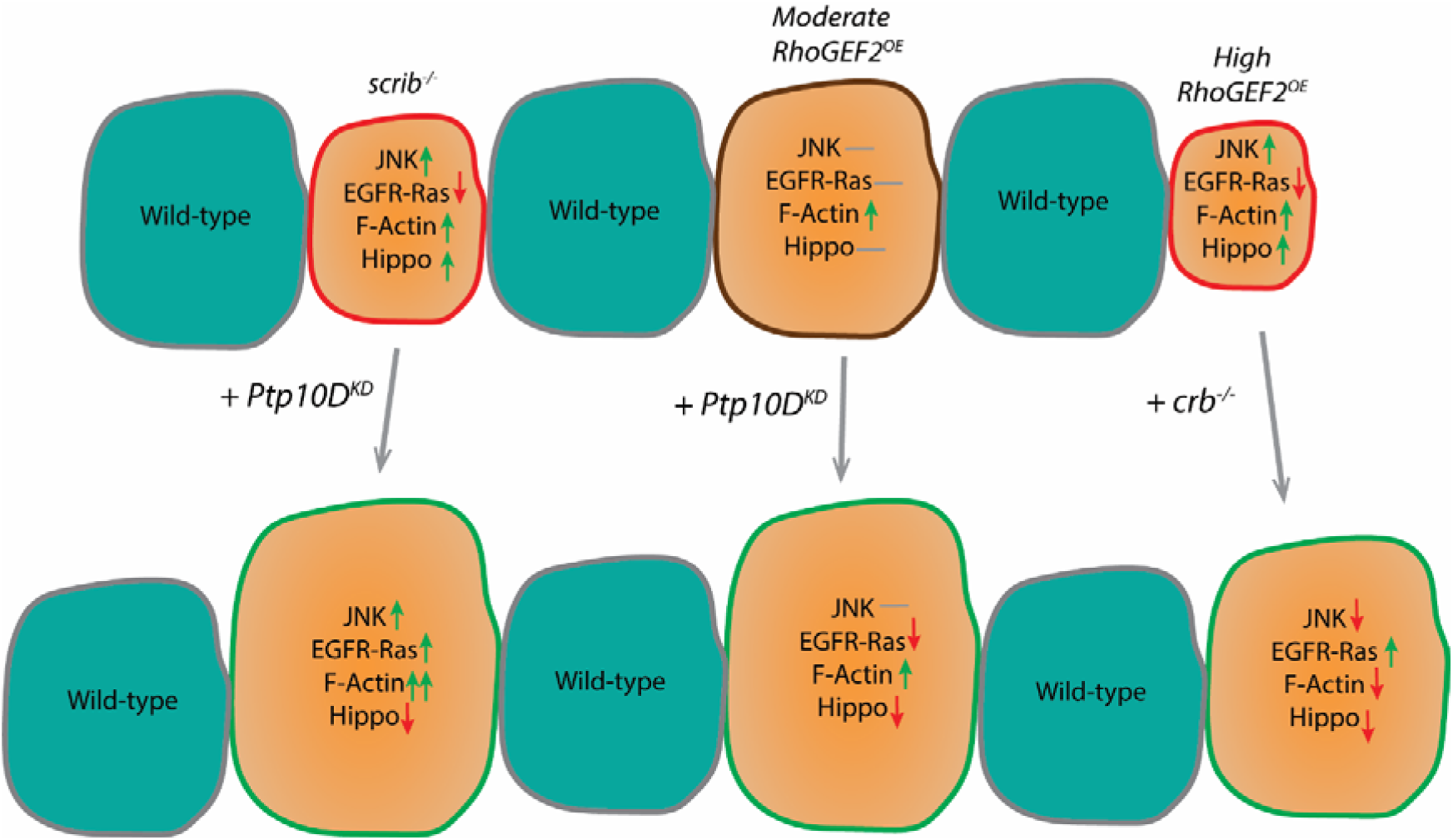
Summary of cell signalling pathway activity upon the different genetic alterations studied in this manuscript. Green arrows indicate upregulation, red arrows indicate downregulation, grey horizontal line indicate no change observed or not investigated. A red clone outline signifies that these clones become a loser clone when surrounded by wild-type clones. A green clone outline indicates that these clones become winner clones relative to wild-type clones, a brown line indicates that no clone size change is observed. Clone sizes are shown to represent relative sizes for each condition. scrib^−/−^ clones are eliminated by cell competition through upregulation of JNK, which is acting through its pro-apoptotic role since EGFR-Ras signalling is blocked by Ptp10D. F-actin is upregulated and so is Hippo pathway signalling. Upon Ptp10D knockdown, EGFR-Ras signalling is upregulated and acts synergistically with JNK resulting in F-actin hyperaccumulation and a reduction of Hippo pathway signalling (Yamamoto, Ohsawa et al. 2017). Moderate RhoGEF2 overexpression clones show elevated active Caspase but are not reduced in size, and show elevated F-actin accumulation, but JNK, EGFR-Ras and Hippo signaling are not affected. Upon Ptp10D knockdown, the Hippo pathway is downregulated, and the EGFR-Ras is slightly downregulated. Given that Hippo signalling is decreased without elevated EGFR-Ras signalling upon Ptp10D knockdown, this indicates that Ptp10D knockdown may be inhibiting the Hippo pathway more directly in moderate RhoGEF2 overexpressing clones to result in increased clone survival. High RhoGEF2 overexpression clones show F-actin accumulation and reduced EGFR-Ras signaling and are eliminated through a JNK dependent mechanism and Hippo upregulation. Upon crb loss, JNK signalling, Hippo signalling and F-actin accumulation are reduced, and EGFR-Ras signalling is upregulated.

Whilst no effect on pERK levels was observed in moderate RhoGEF*2^OE^* clones [Supp Fig 7], high *RhoGEF2^OE^* clones led to pERK levels being diminished compared to *wild-type* [Fig 6D and 6D’’’ (controls in Supp Fig 8); quantified in 6F]. However, upon *crb* loss in *RhoGEF2^OE^* clones, pERK levels were rescued to *wild-type* levels [compare Figs 6D’ and 6D’’’ to 6E’ and 6E’’’; quantified in 6F], indicating that Crb plays a role in inhibiting EGFR-Ras signaling in *RhoGEF2^OE^* clones. Next, we tested whether Crb was involved in F-actin accumulation in *RhoGEF2^OE^*clones. Utilizing Phalloidin staining, we confirmed the previously reported observation that F-Actin accumulates in *RhoGEF2^OE^* clones (Khoo et al., 2013), and furthermore that upon loss of *crb* there was a complete rescue of F-actin accumulation [compare Figs 6G’ and 6H’; quantified in 6I]). Additionally, rescue of the cyst-like phenotype of *RhoGEF2^OE^*clones [compare Figs 6C and 6B – more detailed cyst-like phenotype shown in Fig 2B and C] was observed upon *crb* loss. Taken together, these findings show that *crb* loss in *RhoGEF2^OE^*clones reduces F-actin accumulation and cyst formation.

Importantly, Crb serves as a Hippo pathway regulator, as mutations in *crb* lead to inhibition of the Hippo pathway and the expression of genes that facilitate cell proliferation and inhibit cell death (Robinson et al., 2010). To investigate the impact of *crb* loss on the Hippo pathway in *RhoGEF2^OE^*clones, we analysed expression of the Hippo pathway target Diap1. In *RhoGEF2^OE^* clones alone, Diap1 levels were notably reduced compared to the *wild-type* [Fig 6J’ and 6J’’’ (controls in Supp Fig 8); quantified in Fig 6L], consistent with increased Hippo signaling in *RhoGEF2^OE^*clones. Conversely, and as expected, *RhoGEF2^OE^ crb^−/−^*clones exhibited increased Diap1 staining compared with *RhoGEF2^OE^* clones, indicating decreased Hippo pathway activity [compare Figs 6J’ and 6J’’’ to Fig 6K’ and 6K’’’; quantified in 6L]. Interestingly, in our experiments *crb^−/−^* clones did not elevate Diap1 expression relative to *wild-type* (Supp Fig 8; quantified in Fig 6L), suggesting that in *RhoGEF2^OE^* clones *crb* loss is not simply increasing Diap1 levels through a counteractive effect. Thus, Crb may be playing a specific role in *RhoGEF2^OE^*cell competition by inducing Hippo pathway activity. Altogether, these results show that higher expression of *RhoGEF2^OE^* results in elevated JNK signaling, reduced EGFR-Ras signaling, and elevated Hippo signaling (similar to what occurs in *scrib* mutant clones (Yamamoto et al., 2017)), and that *crb* loss rescues these signaling pathway defects.

### Ptp10D is downregulated in *crb* mutant clones

Finally, we explored whether Crb might be required for Ptp10D protein abundance/membrane localization, since it has been previously reported that it aids in plasma membrane localization of other proteins (Fletcher et al., 2012; Hafezi et al., 2012). We investigated the effect of *crb* loss on Ptp10D protein levels and localization by conducting confocal Z-section analysis of Ptp10D immunostaining in *crb*^−/−^ mosaic eye discs. This examination revealed a notable decrease in Ptp10D levels within *crb^−/−^*eye disc clones, particularly at the apical membrane where it typically localizes [compare Figs 7A and 7B; quantified in 7C]. This result suggests a potential role for Crb in regulating the protein abundance or localization of Ptp10D at the apical membrane. Furthermore, this finding poses an important question of whether Crb is involved in promoting Sas-Ptp10D-mediated cell competition through its effect on Ptp10D abundance/apical membrane localization.

## Discussion

Cell competition is a rapidly expanding research area, but a complex one. In this manuscript, we first explored and expanded upon the findings of previous studies regarding the role of the Ptp10D-Sas axis in polarity-deficient cell elimination (Gerlach et al., 2022; Igaki et al., 2002; Liu et al., 2022; Yamamoto et al., 2017), around which there had been some contention (Gerlach et al., 2022). We also recapitulated observations regarding *RhoGEF2^OE^* clone elimination by cell competition (Brumby et al., 2011; Khoo et al., 2013), and have further demonstrated a role for Sas-Ptp10D signaling in restricting the growth of clones of these cytoskeleton deregulated cells, indicating that loss of polarity is not a prerequisite for Sas-Ptp10D activity. We observed that in cells with moderate *RhoGEF2^OE^*, Ptp10D induced cell competition by activating the Hippo pathway. Additionally, we found that the apical cell polarity protein Crb contributed to the elimination of high *RhoGEF2^OE^* cells in a clonal context, via a mechanism involving normalization of the elevated JNK and Hippo signaling and the reduced EGFR-Ras signaling in high *RhoGEF2^OE^* clones. The results of our findings are summarized in Fig 8. Furthermore, we demonstrated that *crb* mutant cells reduce the expression of and apical membrane localization of Ptp10D, suggesting that Crb plays a key role in Ptp10D signaling during *RhoGEF2^OE^* clone elimination. This study broadens our understanding of Ptp10D signaling in cell competition, illustrating its involvement in eliminating “loser cells” with deregulated actin cytoskeleton.

### Ptp10D’s role in moderate *RhoGEF2^OE^* clones is Hippo dependent but JNK and EGFR independent

In our moderate overexpression model, *RhoGEF2^OE^ Ptp10D^KD^* clones presented significantly higher Diap1 levels compared to *RhoGEF2^OE^ Dicer2*, implying their increased size is due to Hippo pathway inhibition, in line with the canonical *scrib^−/−^ Ptp10D^KD^* model (Yamamoto et al., 2017). Nevertheless, how the Hippo pathway is inactivated in this setting seems to differ, since Ras signaling was not observed to be reduced in moderate *RhoGEF2^OE^* clones or elevated in *RhoGEF2^OE^ Ptp10D^KD^* clones. Since Sas-Ptp10D signaling is known to inhibit EGFR-Ras signaling in *scrib* mutant cells (Yamamoto et al., 2017), it is perplexing why moderate *RhoGEF2^OE^*clones did not also lead to reduced Ras signaling. It is possible that moderate *RhoGEF2^OE^* clones do not strongly activate the Sas-Ptp10D system, or that the potential inhibition of EGFR-Ras signaling by Sas-Ptp10D in this setting might be counteracted by unknown factors in the moderate *RhoGEF2^OE^* clones. Elevated JNK signaling was also not detectable in our moderate *RhoGEF2^OE^* model, in contrast to what occurs in *scrib* mutant clones. F-actin levels are upregulated in moderate *RhoGEF2^OE^* clones but were not significantly affected by *Ptp10D^KD^* in these clones. These findings indicate that in these moderate *RhoGEF2^OE^ Ptp10D^KD^* clones Hippo signaling is being downregulated via a different mechanism to that previously described for *scrib^−/−^ Ptp10D^KD^* clones, where EGFR-Ras and JNK signaling synergize to promote F-actin hyperaccumulation and Hippo pathway inhibition (Yamamoto et al., 2017). It is possible that Ptp10D might be regulating the Hippo pathway directly; for example, since Ptp10D is a phosphatase, it might function to dephosphorylate components of the Hippo pathway, such as Expanded (a Hippo pathway upstream activator). Expanded is phosphorylated by Casien Kinase 1 and then targeted for degradation by the Slimb-β-TrcP ubiquitin ligase (Fulford et al., 2019). If Ptp10D can dephosphorylate Expanded, its knockdown would be expected to lead to higher levels of Expanded phosphorylation and its subsequent degradation, therefore resulting in greater suppression of the Hippo pathway. Alternatively, Ptp10D might regulate signaling pathways that regulate Hippo signaling, such as the Hedgehog pathway (Kagey et al., 2012) or Decapentaplegic pathway (Oh and Irvine, 2011), by dephosphorylation of key components in these pathways.

### *Ptp10D* knockdown rescued the adult eye phenotype of moderate *RhoGEF2^OE^* mosaics

Interestingly, although moderate *RhoGEF2^OE^ Ptp10D^KD^* clones were increased in size within the eye-antennal imaginal discs, the adult eye phenotype was largely rescued and presented with a less severe phenotype than moderate *RhoGEF2^OE^* (*+ Dicer2*) mosaics. This is different to what has been observed so far with loss of the polarity regulators Scrib and Dlg, where *Ptp10D* knockdown increased the severity of the adult eye phenotype (Yamamoto et al., 2017). Furthermore, while no necrotic tissue was found in moderate *RhoGEF2^OE^* or *RhoGEF2^OE^ Ptp10D^KD^*mosaic adult eyes, some of the *RhoGEF2^OE^* adult eye mosaics had protrusions present that were not observed upon *Ptp10D* knockdown. In some *RhoGEF2^OE^* mosaic adult eyes, it seemed like parts of the antenna were present in the eye structure, suggesting that transdetermination events might be occurring, as has been previously observed upon elevation of Rho/Rac signaling (Brumby et al., 2011). Transdetermination has been shown to involve ectopic Wingless signaling (McClure and Schubiger, 2007; Schubiger et al., 2010), so it is possible that *Ptp10D* knockdown might inhibit this pathway. Further work is needed to provide insight into why *Ptp10D*^KD^ rescues the moderate *RhoGEF2^OE^* adult eye phenotype, which will include a more detailed understanding of the signaling pathways regulated by Ptp10D and RhoGEF2.

### Involvement of the apical protein Crb in elimination of *RhoGEF2^OE^*cytoskeletal deregulated clones

Since Crb is a known regulator of the Hippo pathway (reviewed in Richardson and Portela, 2017) and, like Ptp10D, is localized to the apical membrane, we investigated the involvement of *crb* in *RhoGEF2^OE^* clone elimination. We showed that *crb* loss in *RhoGEF2^OE^* clones partially rescued their small size, which indicates that Crb is playing a role in the elimination of *RhoGEF2^OE^* clones. Furthermore, a significant decrease in cell death was observed in *RhoGEF2^OE^ crb*^−/−^ clones when compared to *RhoGEF2^OE^* clones alone. These findings show that Crb plays an important role in the elimination of high overexpression *RhoGEF2* clones.

Surprisingly, loss of *crb* reduced JNK signaling in high *RhoGEF2^OE^*clones, suggesting that JNK upregulation is Crb dependent and, therefore, JNK downregulation might contribute to the increased survival of *crb*^−/−^ *RhoGEF2^OE^* clones. Activation of Rho1, Rok, and Myosin II contribute to JNK pathway activation in *RhoGEF2^OE^* eye-antennal disc clones (Khoo et al., 2013), and Crb might function through this same axis to regulate JNK activity. Furthermore, loss of *crb* in high *RhoGEF2^OE^*clones rescued the low levels of pERK and Diap1 in *RhoGEF2^OE^* clones. These results suggest that the mechanism by which *RhoGEF2^OE^ crb^−/−^* clones increase in size involves elevated EGFR-Ras signaling and reduced Hippo signaling, but since JNK signaling was normalized and F-actin accumulation reduced it is independent of elevated JNK signaling and F-actin accumulation. The reduced F-actin accumulation in *RhoGEF2^OE^ crb^−/−^* clones may be a result of low EGFR-Ras and JNK signaling, since in *scrib^−/−^* clones these pathways act synergistically in promoting F-actin accumulation (Yamamoto et al., 2017). The reduced F-actin accumulation in high *RhoGEF2^OE^ crb^−/−^* clones compared to *RhoGEF2^OE^* clones alone is consistent with these clones being less cyst-like in structure, since these clone structures form upon accumulation of cytoskeletal proteins (Bielmeier et al., 2016). Consistent with this, loss of *crb* in *RhoGEF2^OE^* clones, as in *RhoGEF2^OE^ Ptp10D^KD^* clones, rescued the severe *RhoGEF2^OE^* eye phenotype, which is likely due to deregulation in actomyosin dynamics.

How is Crb functioning to contribute to the elimination of high *RhoGEF2^OE^*clones? It has been previously described that the FERM-binding motif (FBD) of Crb controls actinmyosin dynamics (Flores-Benitez and Knust, 2015), and actinmyosin regulators have been shown to control Hippo signaling (Deng et al., 2015; Fernández et al., 2011; Fletcher et al., 2015; Gaspar et al., 2015; Wong et al., 2015). Therefore, upon *crb* loss in high *RhoGEF2^OE^* clones, a change in regulation of actinmyosin dynamics mediated by Crb might explain the decreased F-actin accumulation observed, as well as contribute to the impairment of the Hippo pathway. Additionally, the FBD of Crb also binds to the Hippo pathway regulator, Expanded, regulating its stability and the activity of the Hippo pathway (Chen et al., 2010; Grzeschik et al., 2010; Robinson et al., 2010), and therefore *crb* loss would be expected to directly result in impaired Hippo signaling in the high *RhoGEF2^OE^* clones, accounting for their increased survival. Consistent with our observations of the importance of Crb in *RhoGEF2 ^OE^* cell competition, we observed an upregulation of Crb in *RhoGEF2^OE^* clones [Supp Fig 9], which might lead to increased Hippo pathway activation.

### Are Crb and Ptp10D acting on the same pathway?

We found that *crb^−/−^* reduced the abundance and apical membrane localization of Ptp10D and therefore hypothesized that the effect observed upon loss of *crb* in *RhoGEF2^OE^* clones might be a direct consequence of deregulated Sas-Ptp10D signaling. The major commonality between *crb* loss and *Ptp10D^KD^*in *RhoGEF2^OE^* clones was the effect on Diap1. Diap1 downregulation in high *RhoGEF2^OE^* clones was rescued to *wild-type* levels by loss of *crb,* while *Ptp10D^KD^* led to increased Diap1 levels in moderate *RhoGEF2^OE^* clones compared to *wild-type* controls. Since loss of *crb* leads to a downregulation of the expression of Ptp10D and its localization to the apical membrane, it is possible that the effect of *crb* loss on the Hippo signaling pathway is dependent, to some extent, on *Ptp10D* downregulation. As mentioned above, Ptp10D may also contribute to the regulation of Expanded activity through its phosphatase activity, preventing the phosphorylation-mediated degradation of Expanded and promoting Hippo activity. Due to technical issues, we were not able to knockdown *Ptp10D* in high *RhoGEF2^OE^* clones and hence could not assess whether *Ptp10D* knockdown phenocopies other effects of *crb* loss in high *RhoGEF2^OE^* clones. Further investigation is needed, involving genetic manipulation to increase RhoGEF2 levels whilst maintaining the same number of *UAS* elements, to determine whether in high *RhoGEF2^OE^* clones, *Ptp10D* knockdown has the same effect as it does in *scrib* mutant clones (Yamamoto et al., 2017) or acts more like *crb* loss in this context.

The Sas-Ptp10D system had previously only been described to have a role in cell competition for the elimination of polarity-deficient cells. Interestingly, here we show that Sas and Ptp10D relocate to the lateral membrane in *RhoGEF2^OE^* clones, as occurs with *scrib^−/−^* clones (Yamamoto et al., 2017), and that *Ptp10D^KD^* in *RhoGEF2^OE^* clones increases *RhoGEF2^OE^* clonal growth. These two findings indicate that Sas-Ptp10D signaling plays a role in restricting the clonal growth of cytoskeletal deregulated cells and, therefore, that polarity-loss is not a prerequisite for Sas-Ptp10D activation. It is possible that in other forms of cell competition where a precise mechanism is unknown, such as in *Stat92E*-overexpressing clones (Rodrigues et al., 2012; Zoranovic et al., 2013), Sas-Ptp10D signaling might also play a role, and it will be important for future endeavours to investigate this question. Additionally, it will be important to determine whether a role for Crb exists in polarity-impaired cell competition, as well as other cell competition scenarios.

## Materials and Methods

### Drosophila stocks

The following fly stocks were used: *FRT82B* (BDSC) #2035, *scrib^1^*(Bilder et al., 2000), *crb^11A22^* (Tepaß and Knust, 1990), *UAS-myr-RFP* (BDSC) #7118, *UAS-Ptp10D-RNAi* (VDRC) #1101, *UAS-Ptp10D-RNAi* (VDRC) #1102, *UAS-sas-RNAi* (VDRC) #39086, *UAS-RhoGEF2* (Padash Barmchi et al., 2005) 3R MARCM stock with eyeless promoter [*ey-FLP, UAS-mCD8-GFP;; tub-GAL4 FRT82B tub-GAL80 / TM6B* (MARCM 82B)] (Lee and Luo, 2001), 3R Reverse MARCM stock with eyeless promoter [*y-, w-, eyFLP2; Act-GAL4, UAS- GFP; tub-GAL80, FRT82B, scrib1 / TM6B*] (Xianjue Ma and JE La Marca). See Supp Table 1 for genotypes of flies used for each figure.

### Husbandry conditions and food recipes

Stocks were maintained at room temperature (RT). Crosses were undertaken at 25°C in an incubator with a 12-hour light/darkness cycle (unless otherwise indicated). Flies were fed with standard semolina-molasses-agar medium supplemented with live yeast or a low protein diet when indicated (See Supp Table 2).

### Immunohistochemistry: Sample preparation and antibodies

L3 eye-antennal discs were dissected in 1X phosphate-buffered saline (PBS) (Amresco #703) and fixed in 4% paraformaldehyde (Alfa Aesar #43368) in PBS with 0.3% Triton X-100 (Amresco #0694) (PBST) for 20-30 min at RT (Note: when using the pERK antibody, tissues were fixed in 8% PFA). After fixation, tissues were washed and permeabilized 3 times for at least 10 minutes in PBS with 0.3% Triton X-100 (PBST), and then blocked for at least 60 minutes at RT in PBST containing 5% bovine serum albumin (BSA) (Sigma-Aldrich #A2153). Unless otherwise indicated, tissues were then incubated overnight at 4°C, in dilutions of primary antibodies in PBST containing 5% BSA. The next day, tissues were washed three times for at least 10 minutes in PBST, then incubated in secondary antibody in PBST containing 5% BSA at RT for 60-120 minutes. Before mounting, tissues were washed in PBST again 3 times for minimum 10 minutes. All samples were mounted in VECTASHIELD® Antifade Mounting Medium with DAPI (#H-1200-10). For cytoskeleton structure visualization (F-actin), Rhodamine Phalloidin fluorescently tagged with RFP was used 1:250 (ThermoFisher Scientific #R415). Utilized primary antibodies: mouse anti-Ptp10D (1:500, DSHB, 8B22F5), rabbit anti-Sas (1:200, Elisabeth Knust), rabbit anti-Cleaved Caspase-3 (1:100, Cell Signaling Technologies #9661), mouse anti-MMP1 (1:100, DSHB, #3B8D12, #5H7B11, and #3A6B4), mouse anti-Diap1 (1:100, Bruce Hay), rabbit anti pERK (1:200, Cell Signaling technologies, #4370), rat anti-Crumbs (1:400, Elisabeth Knust). Utilized secondary antibodies: anti-Mouse IgG, Alexa Fluor 568 (1:500, ThermoFisher Scientific, #A11004), anti-Rabbit IgG, Alexa Fluor 633 (1:500, ThermoFisher Scientific, #A21070), anti-Rat IgG, Alexa Fluor 568 (1:500, ThermoFisher Scientific #A11077).

### Immunohistochemistry: Imaging

Samples were imaged using either a Zeiss LSM 780 or Zeiss LSM 800 laser scanning confocal microscope, and images were processed using Zen 3.0 SR software (Zeiss). Images were generally taken as Z-stacks, but all images shown in this study represent single planes of the Z-stacks, unless otherwise specified. Laser intensity and gain was unchanged within each experimental group.

### Image analysis: Clone volumes

Clone volume quantification was analysed specifically in the eye primordium (antennal primordium was not included). Quantification of tissue volume was analysed with Imaris software (Bitplane). This software allows for the creation of a 3D structure of the eye imaginal disc by combining all the Z-stack images. Using this structure, the volume of the GFP+ tissue could be determined by generating a mask of the GFP+ tissue. Similarly, the total volume of the eye disc was determined by generating a mask of the DAPI+ tissue. Where thresholding was necessary to differentiate between the tissue positive or negative for these different markers, automatic threshold values chosen by Imaris were used so as not to bias the data. Then, to determine the GFP+ tissue to total tissue ratio the following equation was used: (GFP+ volume (μm^3^)/ DAPI+ volume (μm^3^)) *100 = % GFP+ tissue.

### Image analysis: Pixel intensity ratios

To investigate differences in expression levels or activity of different genes/pathways, pixel intensity ratios were determined using FIJI software (Schindelin et al., 2012) were performed. The “mean” pixel intensity of a specific immunostain within the GFP+ clone tissue was measured using the square selection tool and divided by the mean intensity measured in an identical square selection in an adjacent GFP− region. This was repeated 3-5 times per disc, using different clones each time. The number of analysed discs is specified in the relevant figure legends. Clones to be assessed were selected based on their positioning within the tissue, with those near the tissue borders generally avoided due to unusually high accumulations of stains such as for F-actin and Crb. F-actin quantifications were performed in apical z-stacks.

### Image analysis: Spot density ratios

To measure puncta of cleaved Caspase-3, the “Spots” tool in Imaris software was utilized to automatically count the total number of positively stained cells. Thresholding was used to only include spots of 3µm (or larger) in diameter. The “Filter” tool was utilized to automatically quantify the number of positively stained cells specifically localized within GFP+ tissue. To normalize the number of positively stained cells to the amount of tissue, the following calculation was used: number of stained cells in GFP+ tissue / total volume of GFP+ tissue = A, number of stained cells in GFP− tissue / total volume of GFP− tissue = B, and then, A / B = spot density ratio within GFP+ tissue.

### Adult eye imaging and analysis

Unconscious flies were placed in 1.5ml microcentrifuge tubes, and euthanised at −20°C for at least 30 minutes. Flies were then positioned for imaging using a Petri dish with a raised section to allow for consistent positioning of the head. Eyes were then imaged using a “Fully Automated Fluorescence Stereo Microscope Leica M205 FA” and “Leica Application Suite v4.2.0” software (Leica). All flies were imaged the same day they were euthanised to avoid tissue dehydration.

### Eye size quantifications

To analyse for differences in the eye sizes, FIJI software was utilized to measure the area of the eye and the total area of the fly head in pixels. Then, the ratio of the areas using the following equation was determined: Area ratio = Eye area / Head area.

### Adult eye severity phenotyping

To analyse potential increases in the severity of the adult eye phenotype, a “phenotype severity” score was given, where 1 indicates a *wild-type*-like phenotype, 2 indicates a rough eye with some tissue rugosities, 3 indicates a rougher eye with potentially some necrotic speckles, and 4 indicates a strongly rough eye with necrotic tissue or epithelial protrusions.

### Statistical analyses

Statistical analyses were performed, and graphs generated, using Prism 9 (GraphPad). Data from the quantifications of each experiment were plotted in this software, Comparisons between 2 samples were tested for significance using Student’s t-test, while comparisons between 3 or more samples used one-way ANOVAs or Kruskal-Wallis test, with Tukey’s or Dunn’s comparison test, respectively. Normality tests were also used to determine whether the analyses fit the assumptions of each test. A p-value of < 0.05 was used as the cut-off for statistically significant differences.

## Acknowledgements

We thank Peter Burke for help maintaining the *Drosophila* stocks, and all other members of our lab for constructive discussions. We thank the LIMS Bioimaging Facility for microscopy equipment and technical support. We thank the Australian *Drosophila* research community, the BDSC and the VDRC for providing fly stocks, OzDros for stock importation services, and FlyBase for the wealth of information. For antibodies, we thank various laboratories (as indicated in the materials and methods) and the Developmental Studies Hybridoma Bank, which was created by the National Institute of Child Health and Human Development of the National Institutes of Health and is based at The University of Iowa, for supplying antibodies.

## Supplementary Figures

**Supplementary Figure 1.**
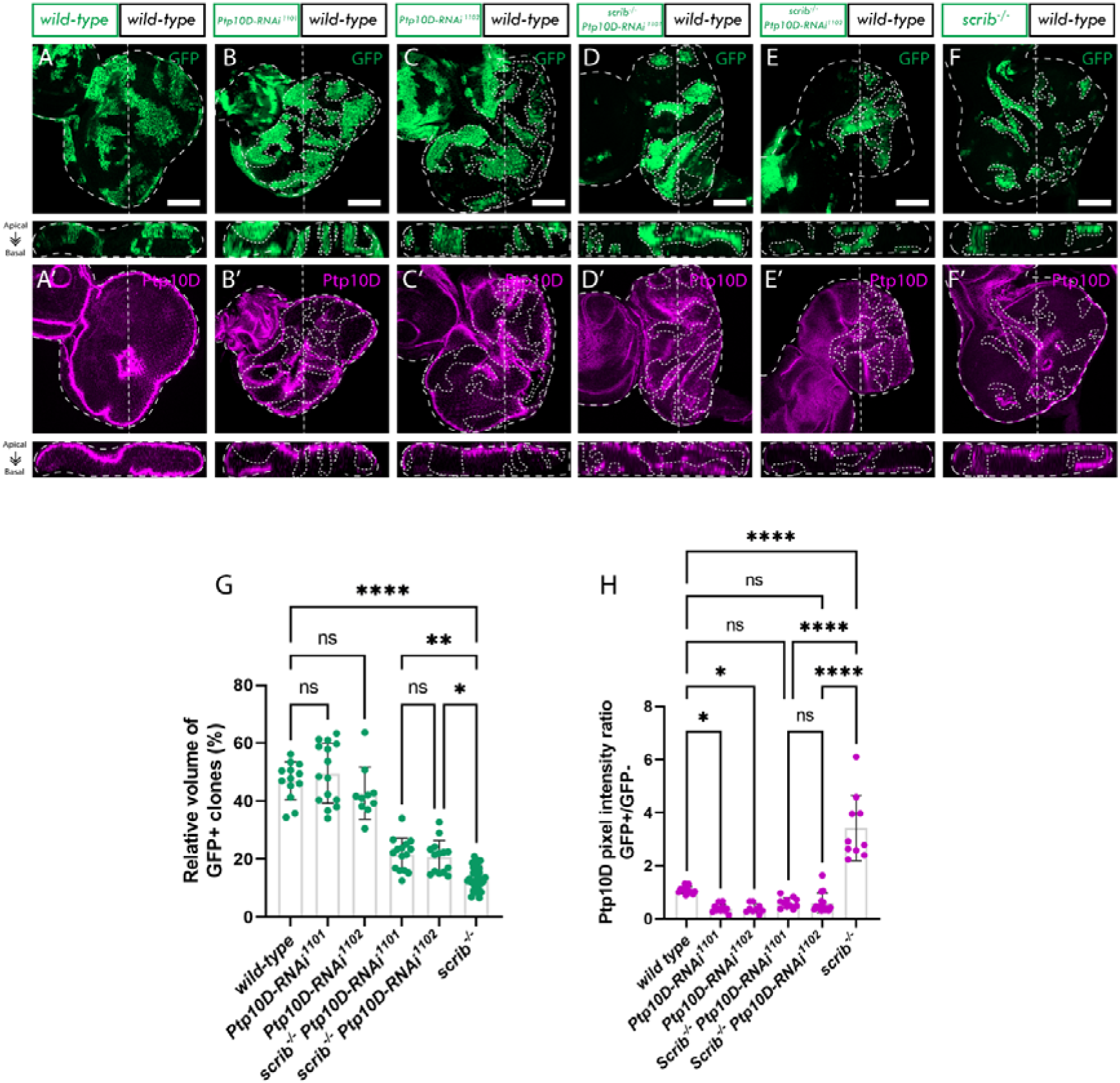
Ptp10D is required for scrib^−/−^ cell elimination. A-F, eye discs of ey-FLP-MARCM-induced mosaics (clones are marked by the presence of GFP and wild-type tissue is unmarked): A) wild-type; B) UAS-Ptp10D-RNAi (VDRC 1101 line); C) UAS-Ptp10D-RNAi (VDRC 1102 line); D) scrib^−/−^ UAS-Ptp10D-RNAi (VDRC 1101 line); E) scrib^−/−^ UAS-Ptp10D-RNAi (VDRC 1102 line); F) scrib^−/−^. A’-F’) Ptp10D immunostains from A-F. G) Quantifications of relative GFP+ clone volume [wild-type (n=13); Ptp10D-RNAi^1101^ (n= 15); Ptp10D-RNAi^1102^ (n=10); scrib^−/−^ Ptp10D-RNAi^1101^ (n=15); scrib^−/−^ Ptp10D-RNAi^1102^ (n=14); scrib^−/−^ (n=32); Statistical test used were one-way ANOVA, p<0.05, with Tukey’s multiple comparison test. In graph: p*: 0.0143; **: 0.0026; ****: p<0.0001]. H) Quantification of Ptp10D immunostain pixel intensity ratio between GFP+ and GFP− clones [wild-type (n=13); Ptp10D-RNAi ^1101^ (n= 11); Ptp10D-RNAi^1102^ (n=9); scrib^−/−^ Ptp10D-RNAi^1101^ (n=11); scrib^−/−^ Ptp10D-RNAi^1102^ (n=15); scrib^−/−^ (n=10); Statistical test used were one-way ANOVA, p<0.05, with Tukey’s multiple comparison test. In graph: *: 0.0188 (wild-type vs Ptp10D-RNAi); *: 0.0325 (wild-type vs Ptp10D-RNAi); ****: p<0.0001]. ns = not significant; error bars = SD; scale bars indicate 50 µm. Images below show an xz cross section of the corresponding eye-antennal disc from the apical (top) to basal (bottom) edge, with the position of the chosen xz sections indicated by a vertical dotted line in the xy images. Note that folding at the edges of the discs can result in Ptp10D being observed basally in the xz sections in these regions, despite the staining still being localized at the apical membrane. Dotted lines surrounding discs or clones illustrate disc/clone boundaries.

**Supplementary Figure 2.**
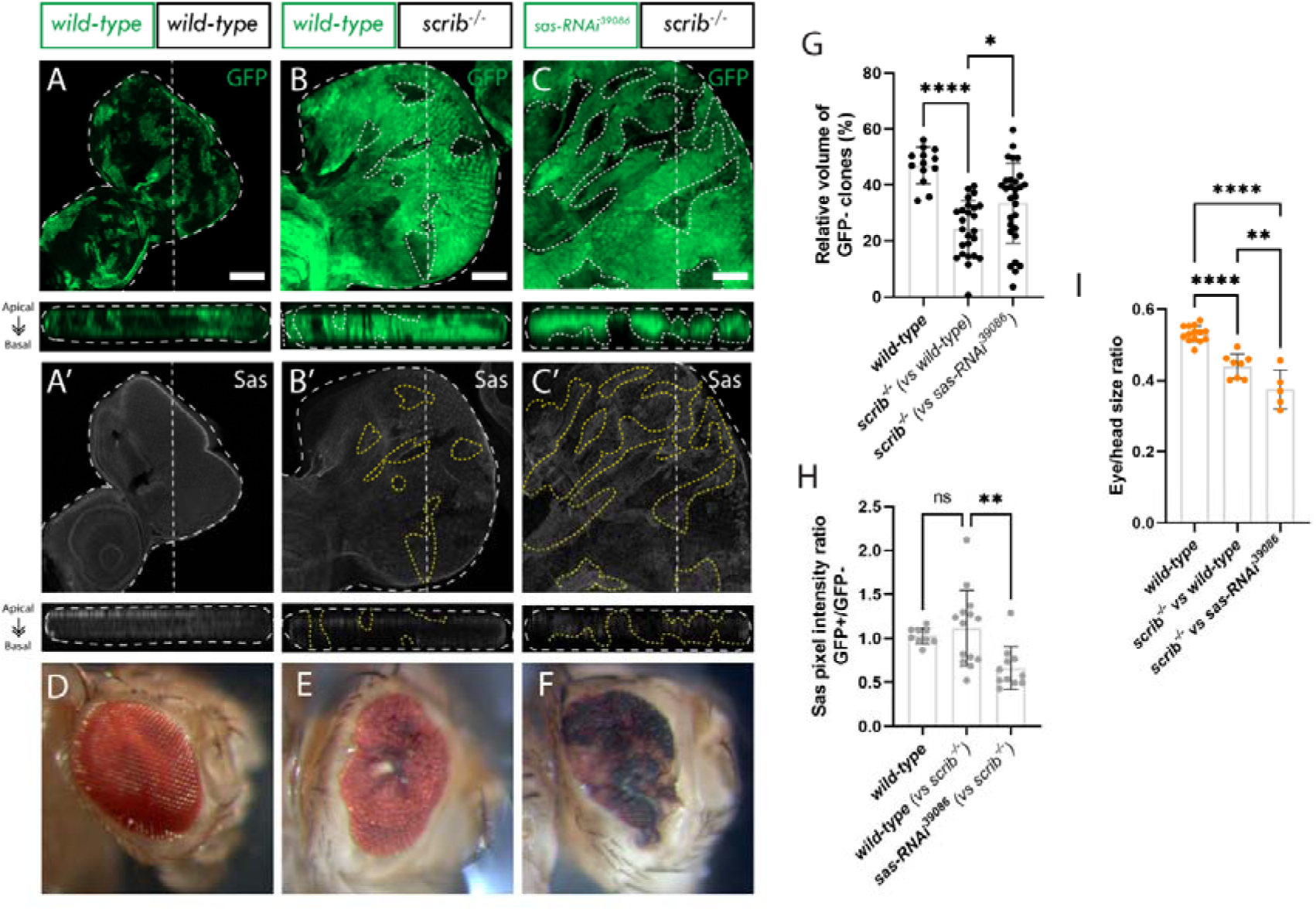
Sas, in wild-type neighbouring cells, is necessary for the elimination of scrib^−/−^ clones in eye disc mosaics. A, eye-antennal discs of the ey-FLP MARCM wild-type mosaic control. B-C, eye-antennal discs of ey-FLP-reverse-MARCM-induced mosaics (clones are marked by the absence of GFP and wild-type tissue marked with GFP): B) scrib^−/−^ (GFP-) and wild-type (GFP+); C) scrib^−/−^ (GFP-) and sas-RNAi^39086^ (GFP+). Note the reverse MARCM stock contains FRT82B scrib^−/−^ Tub-GAL80 and therefore the standard MARCM stock was used as the control (A). The control disc may be smaller relative to the scrib^−/−^ mosaic discs due to scrib^−/−^ clones inducing non-cell autonomous proliferation of the wild-type tissue. A’-C’) Sas immunostains from A - C. D – F) Adult eye mosaics corresponding to above panels; G) Quantification of relative GFP− volume [wild-type vs wild-type (n=13); scrib^−/−^ vs wild-type (n=25); scrib^−/−^ vs sas-RNAi^39086^ (n=29); Statistical test used were one-way ANOVA, with Tukey’s multiple comparison test. In graph: *:0.0149; ****<0.0001]; H) Quantification of Sas pixel intensity ratio between GFP+ and GFP− clones [wild-type vs wild-type (n=9); scrib ^−/−^ vs wild-type (n=14); scrib^−/−^ vs sas-RNAi^39086^ (n=11); Statistical test used were Kruskal-walis test, with Dunn’s multiple comparison test. In graph: **:0.0058]; I) Relative eye to head size quantification [wild-type vs wild-type (n=14); scrib^−/−^ vs wild-type (n=8); scrib^−/−^ vs sas-RNAi^39086^ (n=5); Statistical test used were one-way ANOVA, with Tukey’s multiple comparison test. In graph: **:0.0056; ****<0.0001]. ns = not significant; error bars = SD; scale bars indicate 50 µm. Below each image is an xz cross section of the corresponding eye-antennal disc from the apical (top) to basal (bottom) edge, with the position of the chosen xz sections indicated by a vertical dotted line in the xy images. Dotted lines surrounding discs or clones illustrate disc/clone boundaries.

**Supplementary Figure 3.**
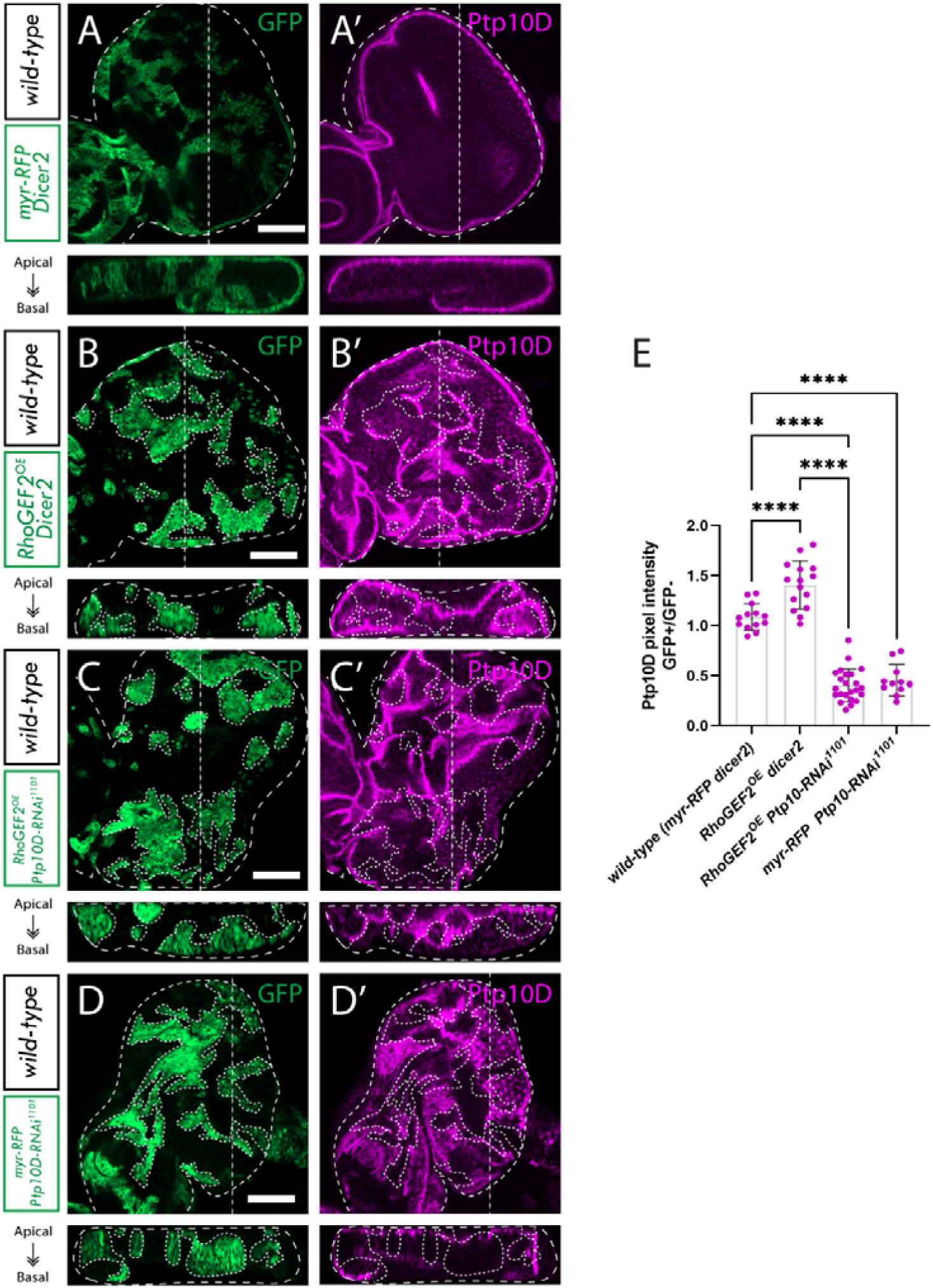
^KD^ increases RhoGEF2^OE^ clonal growth, in RhoGEF2^OE^ eye disc mosaics. A-D, eye discs of ey-FLP-MARCM-induced mosaics (clones are marked by the presence of GFP and wild-type tissue is unmarked): A) wild-type; B) UAS-RhoGEF2^OE^ UAS-Dicer-2; C) UAS-RhoGEF2 UAS-Ptp10D-RNAi (VDRC 1102 line); D) UAS-myr-RFP UAS-Ptp10D-RNAi (VDRC 1101 line). A’-D’) Ptp10D immunostains from A-D. E) Quantification of Ptp10D pixel intensity ratio between GFP+ and GFP− clones [wild-type (n=13); RhoGEF2^OE^ Dicer2 (n=15); RhoGEF2 Ptp10D-RNAi 1101 (n=23); myr-RFP UAS-Ptp10D-RNAi 1101 (n=11); Statistical test used were one-way ANOVA, p<0.05, with Tukey’s multiple comparison test. In graph: ****p <0.0001]. ns = not significant; error bars = SD; scale bars indicate 50 µm. Below each image is an xz cross section of the corresponding eye-antennal disc from the apical (top) to basal (bottom) edge, with the position of the chosen xz sections indicated by a vertical dotted line in the xy images. Note that folding at the edges of the discs can result in Ptp10D being observed basally in the xz sections in these regions, despite the staining still being localized at the apical membrane. Dotted lines surrounding discs or clones illustrate disc/clone boundaries.

**Supplementary Figure 4.**
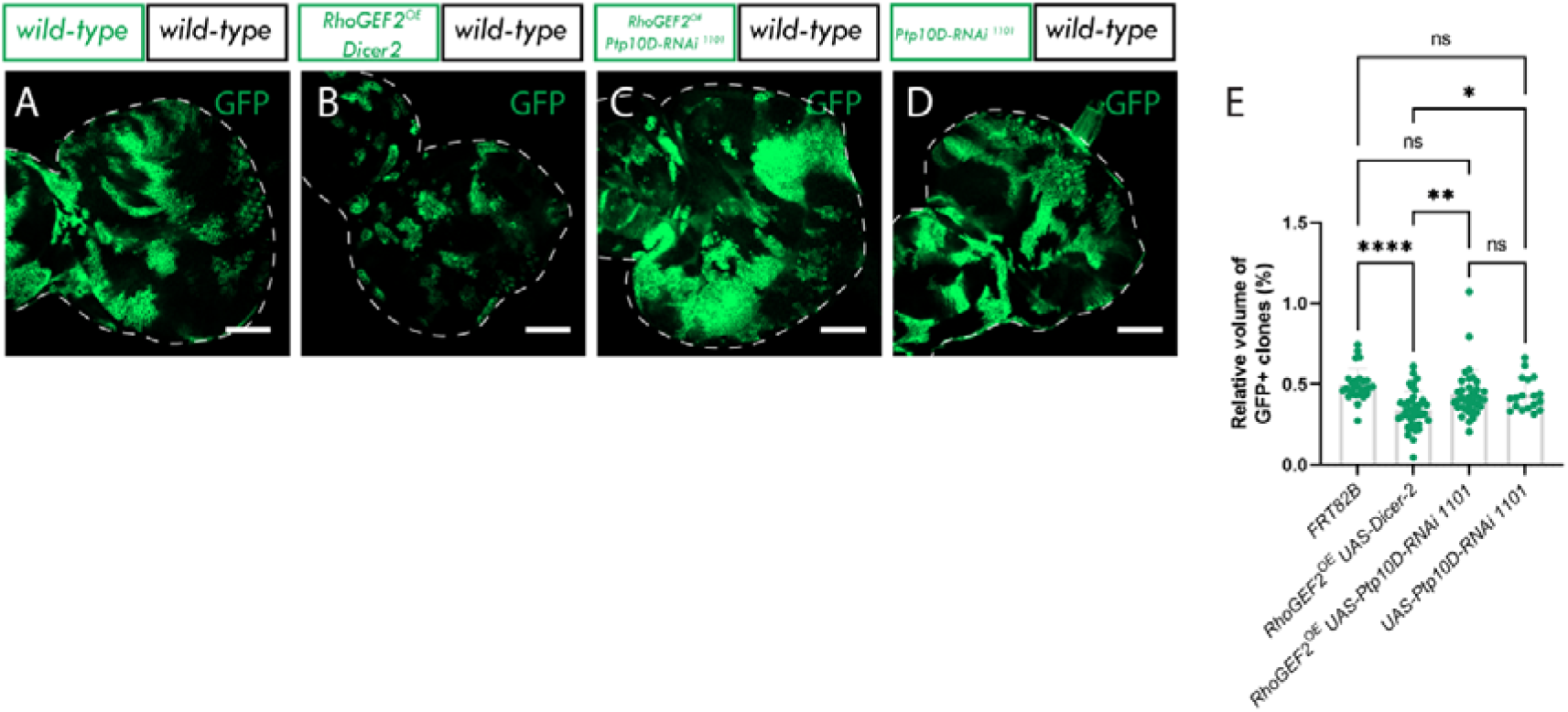
Moderate overexpressing RhoGEF2 clones are eliminated by cell-competition in a low-protein diet. A-D, eye discs of ey-FLP-MARCM-induced mosaics (clones are marked by the presence of GFP and wild-type tissue is unmarked): A) wild-type; B) UAS-RhoGEF2^OE^ UAS-Dicer-2; C) UAS-RhoGEF2 UAS-Ptp10D-RNAi (VDRC 1102 line); D) UAS-Ptp10D-RNAi (VDRC 1101 line). E) Quantification of the relative GFP+ clone volume [wild-type (n=27); RhoGEF2^OE^ Dicer2 (n=40); RhoGEF2 Ptp10D-RNAi 1101 (n=35); myr-RFP UAS-Ptp10D-RNAi 1101 (n=18); Statistical test used were one-way ANOVA, p<0.05, with Tukey’s multiple comparison test. In graph: *: 0.0341; **: 0.0035; ****p <0.0001]. ns = not significant; error bars = SD; scale bars indicate 50 µm. Dotted lines surrounding discs indicate disc boundaries.

**Supplementary Figure 5.**
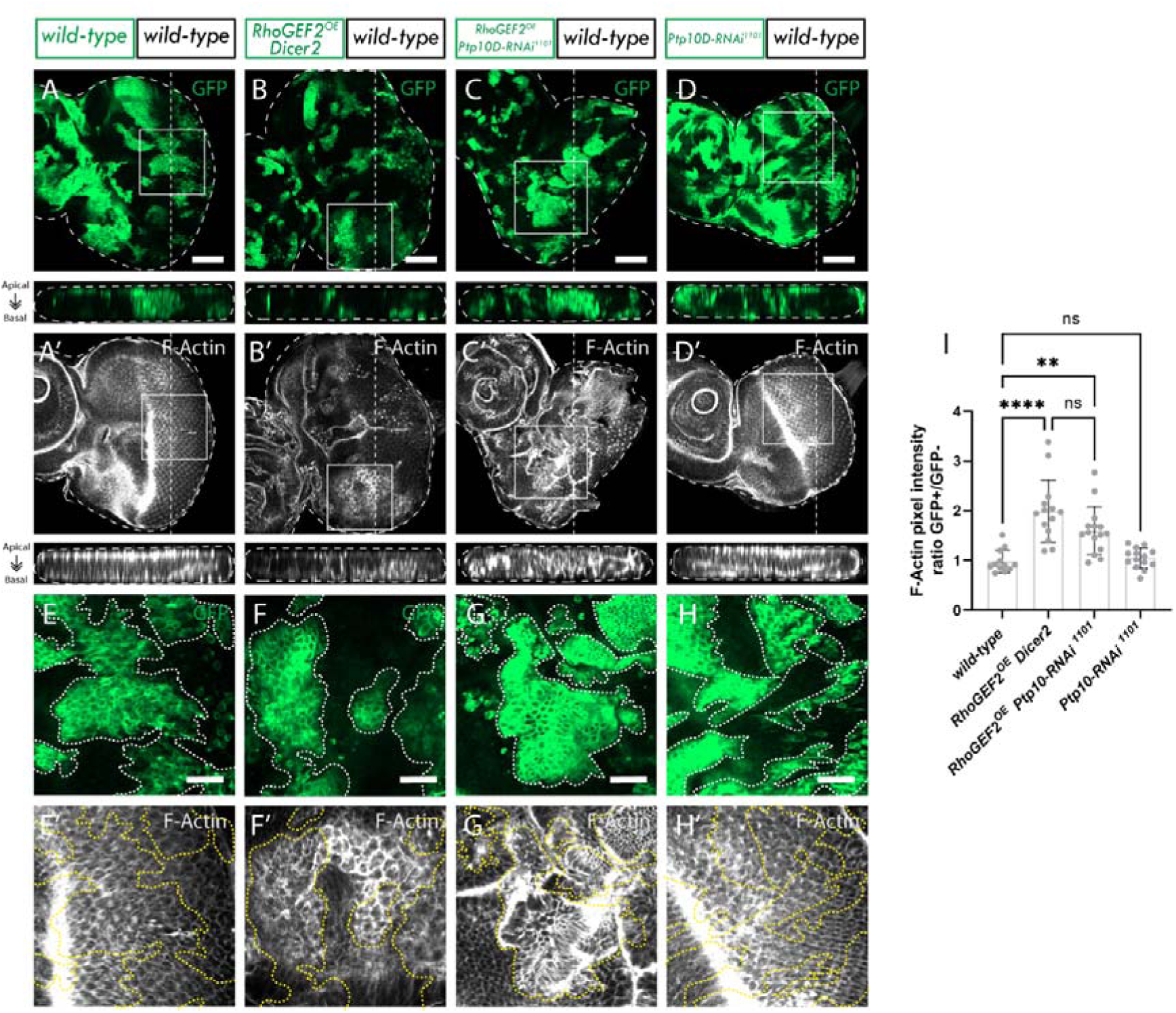
Knockdown of Ptp10D in RhoGEF2^OE^ clones does not significantly affect the increased apical F-actin accumulation in RhoGEF2^OE^ clones, in eye disc mosaic clones. A-D, eye discs of ey-FLP-MARCM-induced mosaics (clones are marked by the presence of GFP and wild-type tissue is unmarked): A) wild-type; B) UAS-RhoGEF2^OE^ UAS-Dicer-2; C) UAS-RhoGEF2 UAS-Ptp10D-RNAi (VDRC 1102 line); D) UAS-myr-RFP UAS-Ptp10D-RNAi (VDRC 1101 line); A’-D’) F-Actin stains from A-D. White squares indicate magnified sections shown below. E-H) Magnified sections from A-D. E’-F’) F-Actin immunostains from E-H; I) Quantification of F-Actin pixel intensity ratio between GFP+ and GFP− clones [wild-type (n=12); RhoGEF2^OE^ Dicer2 (n=14); RhoGEF2 Ptp10D-RNAi 1101 (n=15); myr-RFP UAS-Ptp10D-RNAi 1101 (n=14); Statistical test used were one-way ANOVA, p<0.05, with Tukey’s multiple comparison test. In graph: **0.0063; ****<0.0001]; ns = not significant; error bars = SD; scale bars in A to D indicate 50 µm; scale bars in E to H indicate 20 µm. Dotted lines surrounding discs or clones illustrate disc/clone boundaries.

**Supplementary Figure 6.**
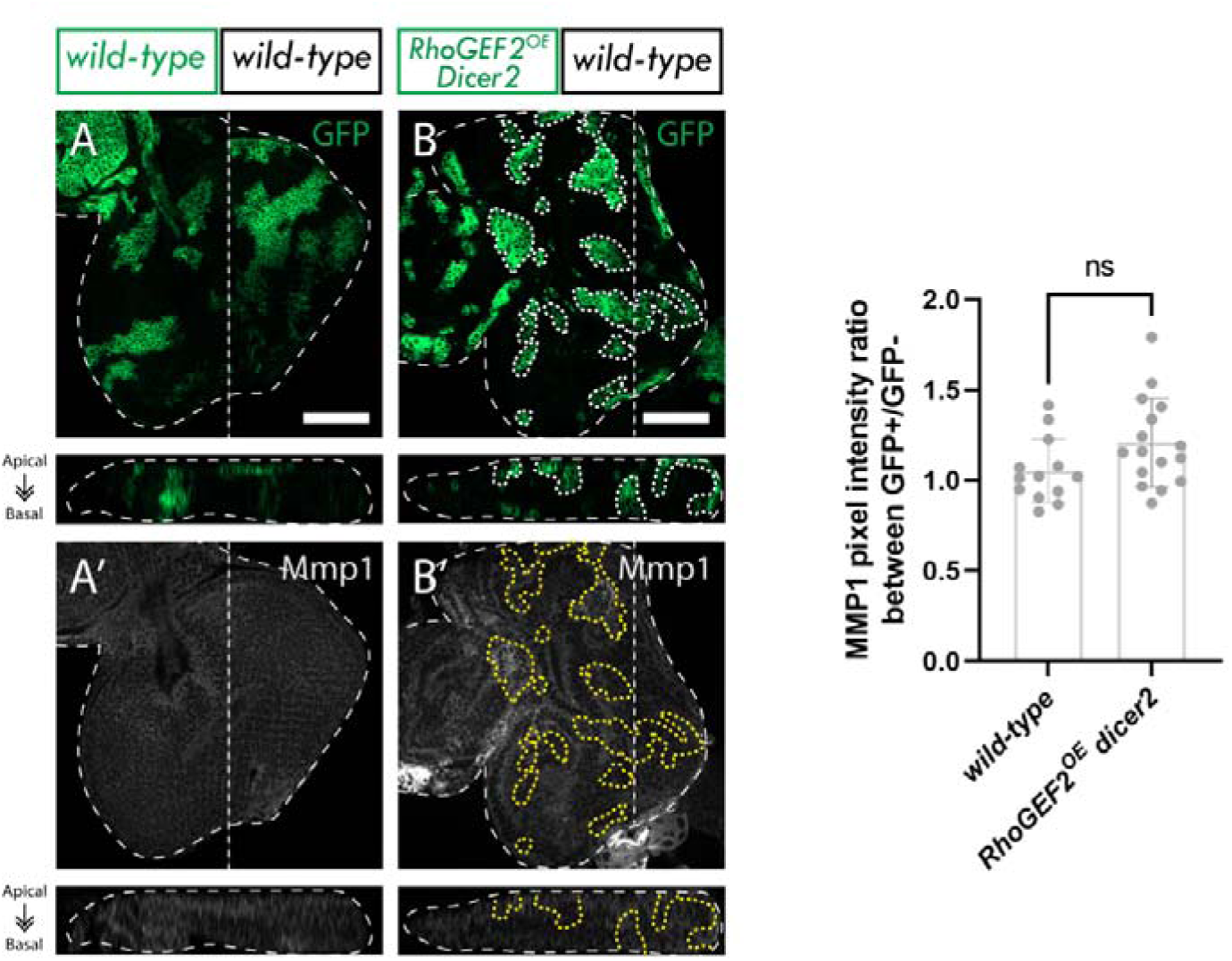
JNK signalling is not activated by moderate overexpression of RhoGEF2. A-B, eye discs of ey-FLP-MARCM-induced mosaics (clones are marked by the presence of GFP and wild-type tissue is unmarked): A) wild-type; B) UAS-RhoGEF2^OE^ UAS-Dicer-2. A’-B’) Mmp1 immunostains from A-B. C) Quantification of MMP1 immunostain pixel intensity ratio between GFP+ and GFP− clones [wild-type (n=13); RhoGEF2^OE^ dicer2 (n=16); Statistical test used was unpaired t-test, p<0.05. p=0.0646]. ns = not significant; error bars = SD; Scale bars indicate 50 µm. Below each image is an xz cross section of the corresponding eye-antennal disc from the apical (top) to basal (bottom) edge, with the position of the chosen xz sections indicated by a vertical dotted line in the xy images. Dotted lines surrounding discs or clones illustrate disc/clone boundaries.

**Supplementary Figure 7.**
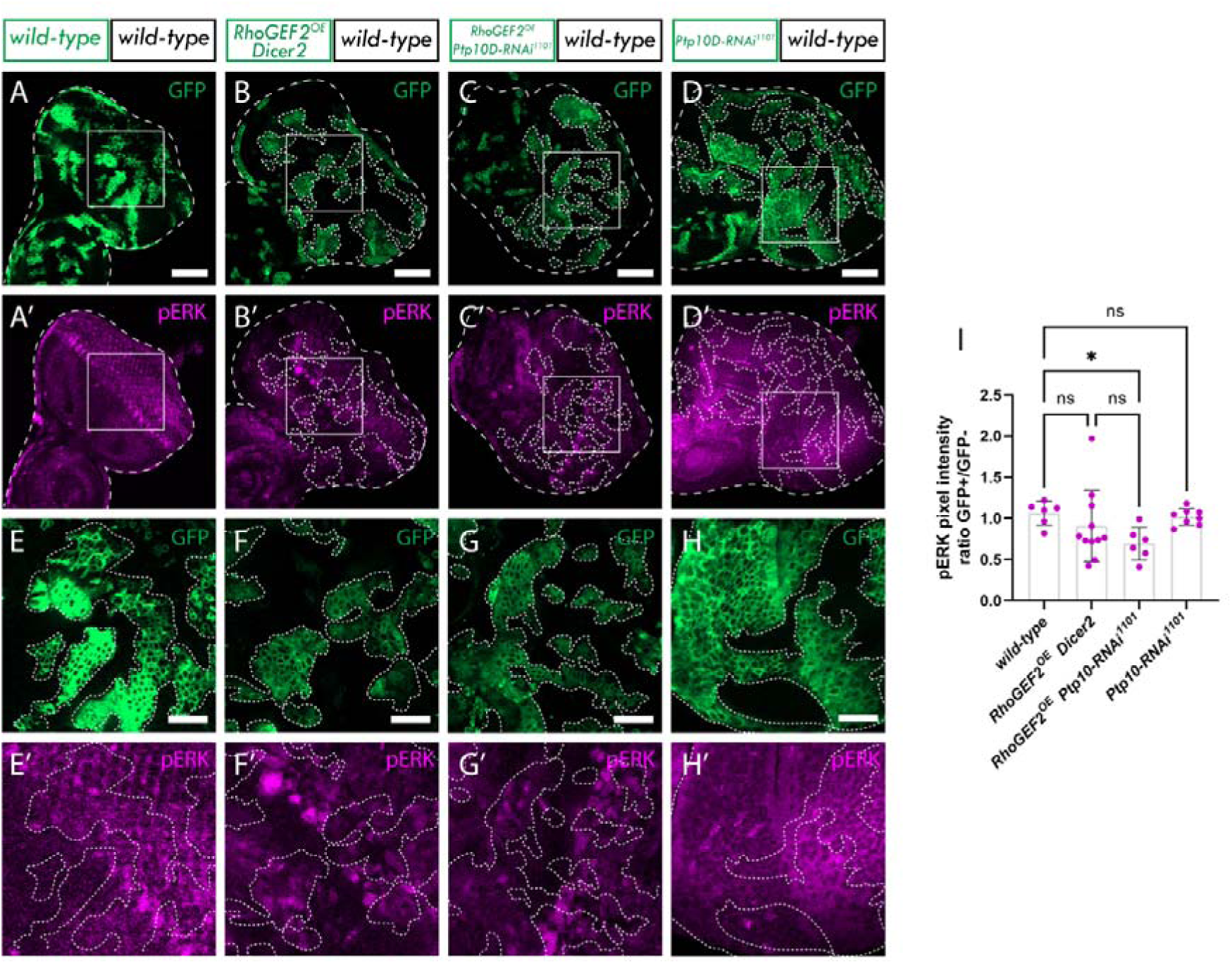
Ptp10D knockdown does not increase Ras signalling in RhoGEF2^OE^ clones. A-D, eye discs of ey-FLP-MARCM-induced mosaics (clones are marked by the presence of GFP and wild-type tissue is unmarked): A) wild-type; B) UAS-RhoGEF2^OE^ UAS-Dicer-2; C) UAS-RhoGEF2 UAS-Ptp10D-RNAi (VDRC 1102 line); D) UAS-myr-RFP UAS-Ptp10D-RNAi (VDRC 1101 line); A’-D’) pERK immunostains from A-D. White squares indicate magnified sections shown below. E-H) Magnified sections from A-D. E’-F’) pERK immunostains from E-F. I) Quantification of pERK pixel intensity ratio between GFP+ and GFP− clones [wild-type (n=6); RhoGEF2^OE^ Dicer2 (n=11); RhoGEF2 Ptp10D-RNAi 1101 (n=6); myr-RFP UAS-Ptp10D-RNAi 1101 (n=8); Statistical test used were Kruskal-Wallis test, p<0.05, with Dunn’s multiple comparison test. In graph: *0.0418]; ns = not significant; error bars = SD; scale bars in A to D indicate 50 µm; scale bars in E to H indicate 20 µm. Dotted lines surrounding discs or clones illustrate disc/clone boundaries.

**Supplementary Figure 8.**
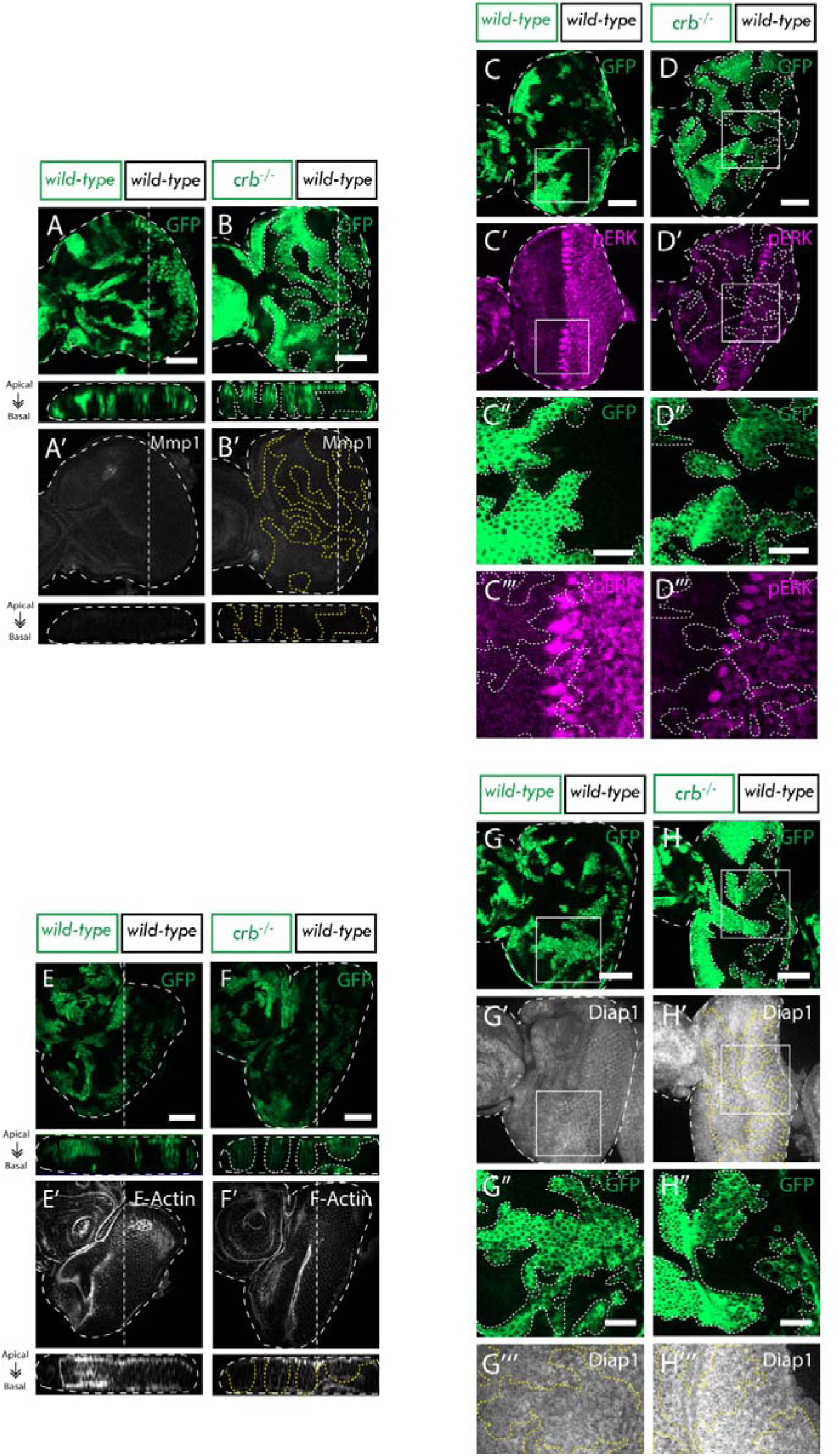
Controls for Figure 6. crb loss decreases apical F-actin accumulation, rescues the elevated JNK signalling, rescues Ras signalling downregulation and reduces the elevated Hippo signalling in RhoGEF2^OE^ clones. A-B, Eye discs of ey-FLP-MARCM-induced mosaics (clones are marked by the presence of GFP and wild-type tissue is unmarked): A) wild-type; B) crb^−/−^; A’-B’) Mmp1 stains from A-B. C-D, Eye discs of ey-FLP-MARCM-induced mosaics (clones are marked by the presence of GFP and wild-type tissue is unmarked): C) wild-type D) crb^−/−^; C’-D’) pERK immunostains from C-D. White squares indicate magnified sections shown in panels below; C’’-D’’) Magnified sections from C-D; C’’’-D’’’) pERK imunostains from C’’-D’’. E-F, Eye discs of ey-FLP-MARCM-induced mosaics (clones are marked by the presence of GFP and wild-type tissue is unmarked): E) wild-type; F) crb^−/−^; E’-F’) F-Actin stains from E-F. G-H, Eye discs of ey-FLP-MARCM-induced mosaics (clones are marked by the presence of GFP and wild-type tissue is unmarked): G) wild-type H) crb^−/−^; G’-H’) pERK immunostains from G-H. White squares indicate magnified sections shown in panels below; G’’-H’’) show magnified sections from G-H; G’’’-H’’’) Diap1 stains in G’’-H’’. ns = not significant; error bars = SD; A, B, C, D, E, F, G, H scale bars indicate 50 µm. C’’, D’’, G’’, H’’) scale bars indicate 20 µm. In A, A’, B, B’, E, E’, F and F’, images below show an xz cross section of the corresponding eye-antennal disc from the apical (top) to basal (bottom) edge, with the position of the chosen xz sections indicated by a vertical dotted line in the xy images. Dotted lines surrounding discs or clones illustrate disc/clone boundaries.

**Supplementary Figure 9.**
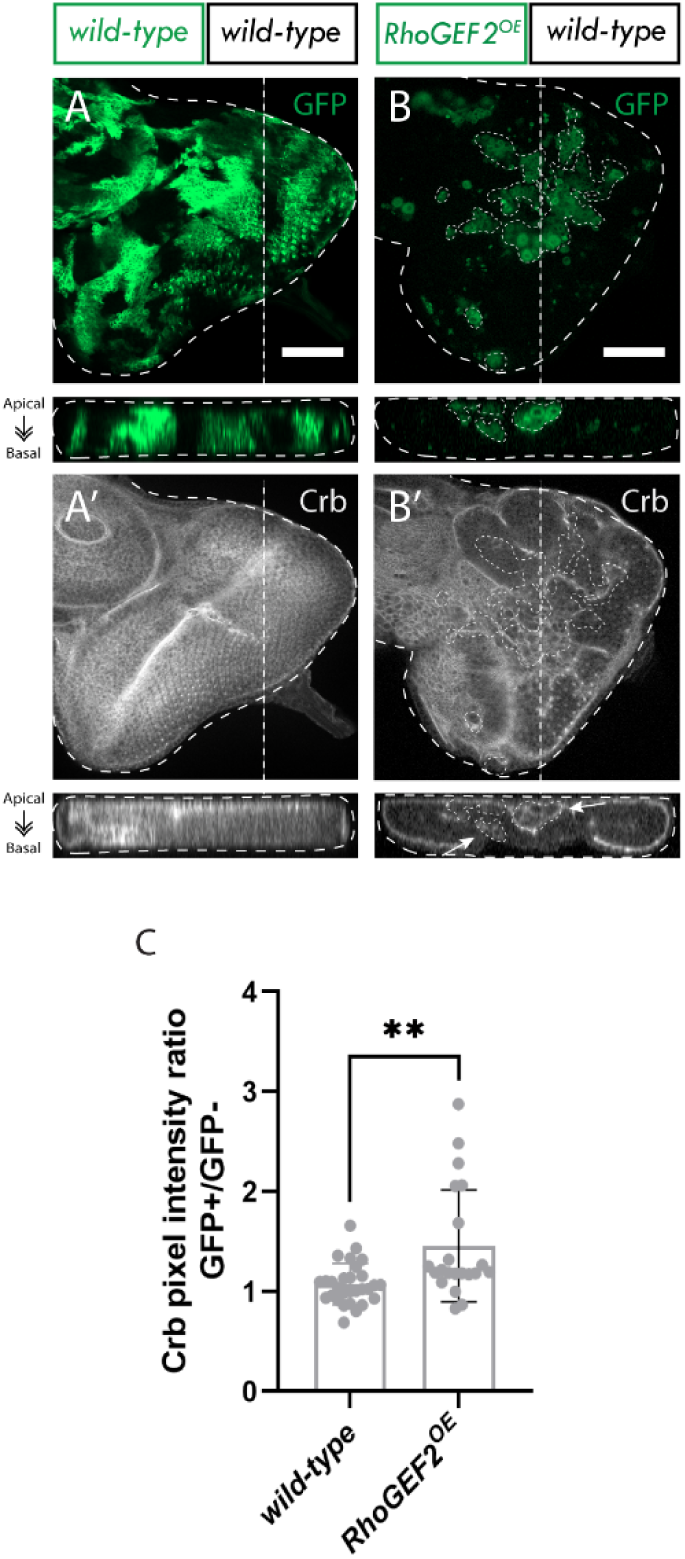
Crb accumulates in RhoGEF2^OE^ clones. A-B, eye discs of ey-FLP-MARCM-induced mosaics (clones are marked by the presence of GFP and wild-type tissue is unmarked): A) wild-type; B) RhoGEF2^OE^. A’ - B’) Crb immunostains. Arrows indicate Crb accumulation in GFP+ clones. C) Quantification of Crb pixel intensity ratio between GFP+ and GFP− clones [wild-type (n=21); RhoGEF2^OE^ (n=29); Statistical test used was Mann Whitney Test. In graph: *P-value: 0.0013]. ns = not significant; error bars = SD; scale bars indicate 50 µm. Below each image is an xz cross section of the corresponding eye-antennal disc from the apical (top) to basal (bottom) edge, with the position of the chosen xz sections indicated by a vertical dotted line in the xy images. Note that folding at the edges of the discs can result in Crb being observed basally in the xz sections in these regions, despite the staining still being localized at the apical membrane. Dotted lines surrounding discs or clones illustrate disc/clone boundaries.

