## Supplementary Tables for "Actin cytoskeletal deregulation, caused by RhoGEF2 overexpression, induces cell competition dependent on Ptp10D, Crumbs, and the Hippo signaling pathway"

**Supp Table 1. Genotypes**

**Fig 1**

A,F,J,O: *eyFLP, UAS-mCD8-GFP /+ ;; tub-GAL4, FRT82B, tub-GAL80/+*

B,G,M,R: *eyFLP, UAS-mCD8-GFP /+ ;; tub-GAL4, FRT82B, tub-GAL80/ FRT82B, scrib^1^*

C,H: *eyFLP, UAS-mCD8-GFP /+ ;; tub-GAL4, FRT82B, tub-GAL80/ FRT82B, scrib^1^, UAS-Ptp10D-RNAi^1101^*

D,I: *eyFLP, UAS-mCD8-GFP /+; UAS-Ptp10D-RNAi^1102^ / + ; tub-GAL4, FRT82B, tub-GAL80/ FRT82B, scrib^1^*

*K,P: eyFLP, UAS-mCD8-GFP /+;* *UAS-Ptp10D UAS.cTa / + ; tub-GAL4, FRT82B, tub-GAL80/ FRT82B*

*L,Q: eyFLP, UAS-mCD8-GFP /+; UAS-Ptp10D UAS.cTa /+ ; tub-GAL4, FRT82B, tub-GAL80/FRT82B, scrib^1^*

**Fig 2**

A, G: *eyFLP, UAS-mCD8-GFP /+ ;; tub-GAL4, FRT82B, tub-GAL80/+*

B, C, H: *eyFLP, UAS-mCD8-GFP /+ ; UAS-RhoGEF2/ +; tub-GAL4, FRT82B, tub-GAL80/FRT82B*

**Fig 3**

A, E: *eyFLP, UAS-mCD8-GFP /+ ;; tub-GAL4, FRT82B, tub-GAL80/+*

B, F: *eyFLP, UAS-mCD8-GFP /+ ; UAS-RhoGEF2/+; tub-GAL4, FRT82B, tub-GAL80/UAS- Dicer2*

C, G: *eyFLP, UAS-mCD8-GFP /+ ; UAS-RhoGEF2/+; tub-GAL4, FRT82B, tub-GAL80/UAS-Ptp10D-RNAi^1101^*

D, H: *eyFLP, UAS-mCD8-GFP /+ ;myr-RFP/+; tub-GAL4, FRT82B, tub-GAL80/UAS-Ptp10D-RNAi^1101^*

**Fig 4**

A, E: *eyFLP, UAS-mCD8-GFP /+ ;; tub-GAL4, FRT82B, tub-GAL80/+*

B, F: *eyFLP, UAS-mCD8-GFP /+ ; UAS-RhoGEF2/+; tub-GAL4, FRT82B, tub-GAL80/UAS- Dicer2*

C, G: *eyFLP, UAS-mCD8-GFP /+ ; UAS-RhoGEF2/+; tub-GAL4, FRT82B, tub-GAL80/UAS-Ptp10D-RNAi^1101^*

D, H: *eyFLP, UAS-mCD8-GFP /+ ;; tub-GAL4, FRT82B, tub-GAL80/UAS-Ptp10D-RNAi^1101^*

**Fig 5**

A, E: *eyFLP, UAS-mCD8-GFP /+ ;; tub-GAL4, FRT82B, tub-GAL80/+*

B, F: *eyFLP, UAS-mCD8-GFP /+ ; UAS-RhoGEF2/ +; tub-GAL4, FRT82B, tub-GAL80/FRT82B*

C, G: *eyFLP, UAS-mCD8-GFP /+ ; UAS-RhoGEF2/ +; tub-GAL4, FRT82B, tub-GAL80/FRT82B, crb^11A22^*

D, H: *eyFLP, UAS-mCD8-GFP /+ ;; tub-GAL4, FRT82B, tub-GAL80/FRT82B, crb^11A22^*

**Fig 6**

*A,D,G,J: B, F: eyFLP, UAS-mCD8-GFP /+ ; UAS-RhoGEF2/ +; tub-GAL4, FRT82B, tub-GAL80/FRT82B*

*B,E,H,K: eyFLP, UAS-mCD8-GFP /+ ; UAS-RhoGEF2/ +; tub-GAL4, FRT82B, tub-GAL80/FRT82B, crb^11A22^*

**Fig 7**

A: *eyFLP, UAS-mCD8-GFP /+ ;; tub-GAL4, FRT82B, tub-GAL80/+*

*B: eyFLP, UAS-mCD8-GFP /+ ;; tub-GAL4, FRT82B, tub-GAL80/FRT82B, crb^11A22^*

**Supp Fig 1**

A: *eyFLP, UAS-mCD8-GFP /+ ;; tub-GAL4, FRT82B, tub-GAL80/+*

B: *eyFLP, UAS-mCD8-GFP /+ ;; tub-GAL4, FRT82B, tub-GAL80/ FRT82B, UAS-Ptp10D-RNAi^1101^*

C: *eyFLP, UAS-mCD8-GFP /+; UAS-Ptp10D-RNAi^1102^ / + ; tub-GAL4, FRT82B, tub-GAL80/ FRT82B*

D: *eyFLP, UAS-mCD8-GFP /+ ;; tub-GAL4, FRT82B, tub-GAL80/ FRT82B, scrib^1^, UAS-Ptp10D-RNAi^1101^*

E: *eyFLP, UAS-mCD8-GFP /+; UAS-Ptp10D-RNAi^1102^ / + ; tub-GAL4, FRT82B, tub-GAL80/ FRT82B, scrib^1^*

F: *eyFLP, UAS-mCD8-GFP /+ ;; tub-GAL4, FRT82B, tub-GAL80/ FRT82B, scrib^1^*

**Supp Fig 2**

*A, D: eyFLP, UAS-mCD8-GFP /+ ;; tub-GAL4, FRT82B, tub-GAL80/+*

*B, E: y-w-, eyFLP2/+; Act>GAL4, UAS-GFP/+; tubGAL80, FRT82B, scrib1/FRT82B*

*C, F: y-w-, eyFLP2/+; Act>GAL4, UAS-GFP/+; tubGAL80, FRT82B, scrib1/FRT82B, UAS-sas-RNAi^39086^*

**Supp Fig 3**

A: *eyFLP, UAS-mCD8-GFP /+ ;; tub-GAL4, FRT82B, tub-GAL80/+*

B: *eyFLP, UAS-mCD8-GFP /+ ; UAS-RhoGEF2/+; tub-GAL4, FRT82B, tub-GAL80/UAS- Dicer2*

C: *eyFLP, UAS-mCD8-GFP /+ ; UAS-RhoGEF2/+; tub-GAL4, FRT82B, tub-GAL80/UAS-Ptp10D-RNAi^1101^*

D: *eyFLP, UAS-mCD8-GFP /+ ;; tub-GAL4, FRT82B, tub-GAL80/UAS-Ptp10D-RNAi^1101^*

**Supp Fig 4**

A: *eyFLP, UAS-mCD8-GFP /+ ;; tub-GAL4, FRT82B, tub-GAL80/+*

B: *eyFLP, UAS-mCD8-GFP /+ ; UAS-RhoGEF2/+; tub-GAL4, FRT82B, tub-GAL80/UAS- Dicer2*

C: *eyFLP, UAS-mCD8-GFP /+ ; UAS-RhoGEF2/+; tub-GAL4, FRT82B, tub-GAL80/UAS-Ptp10D-RNAi^1101^*

D: *eyFLP, UAS-mCD8-GFP /+ ;; tub-GAL4, FRT82B, tub-GAL80/UAS-Ptp10D-RNAi^1101^*

**Supp Fig 5**

A, E: *eyFLP, UAS-mCD8-GFP /+ ;; tub-GAL4, FRT82B, tub-GAL80/+*

B, F: *eyFLP, UAS-mCD8-GFP /+ ; UAS-RhoGEF2/+; tub-GAL4, FRT82B, tub-GAL80/UAS- Dicer2*

C, G: *eyFLP, UAS-mCD8-GFP /+ ; UAS-RhoGEF2/+; tub-GAL4, FRT82B, tub-GAL80/UAS-Ptp10D-RNAi^1101^*

D, H: *eyFLP, UAS-mCD8-GFP /+ ;; tub-GAL4, FRT82B, tub-GAL80/UAS-Ptp10D-RNAi^1101^*

**Supp Fig 6**

A: *eyFLP, UAS-mCD8-GFP /+ ;; tub-GAL4, FRT82B, tub-GAL80/+*

B: *eyFLP, UAS-mCD8-GFP /+ ; UAS-RhoGEF2/+; tub-GAL4, FRT82B, tub-GAL80/UAS- Dicer2*

**Supp Fig 7**

A, E: *eyFLP, UAS-mCD8-GFP /+ ;; tub-GAL4, FRT82B, tub-GAL80/+*

B, F: *eyFLP, UAS-mCD8-GFP /+ ; UAS-RhoGEF2/+; tub-GAL4, FRT82B, tub-GAL80/UAS- Dicer2*

C, G: *eyFLP, UAS-mCD8-GFP /+ ; UAS-RhoGEF2/+; tub-GAL4, FRT82B, tub-GAL80/UAS-Ptp10D-RNAi^1101^*

D, H: *eyFLP, UAS-mCD8-GFP /+ ;; tub-GAL4, FRT82B, tub-GAL80/UAS-Ptp10D-RNAi^1101^*

**Supp Fig 8**

*A,C,E,G: eyFLP, UAS-mCD8-GFP /+ ;; tub-GAL4, FRT82B, tub-GAL80/+*

*B,D,F,H: eyFLP, UAS-mCD8-GFP /+ ;; tub-GAL4, FRT82B, tub-GAL80/FRT82B, crb^11A22^*

**Supp Fig 9**

A, E: *eyFLP, UAS-mCD8-GFP /+ ;; tub-GAL4, FRT82B, tub-GAL80/+*

B, F: *eyFLP, UAS-mCD8-GFP /+ ; UAS-RhoGEF2/ +; tub-GAL4, FRT82B, tub-GAL80/FRT82B*

Supp Table 2. *Drosophila* food recipes

| Reagents | Molasses based food recipe  (g/L) | /Low protein diet  (g/L) |
| --- | --- | --- |
| Molasses* | 93 | - |
| Yeast | 60 | 10.7 |
| Agar | 5 | 4.8 |
| Glucose (Dextrose) | 10.6 | 47.6 |
| Sugar (Raw) | - | 23.8 |
| Semolina | 88 | 59.5 |
| Potassium Sodium Tartrate Tetrahydrate | - | 7.14 |
| Calcium Chloride Dihydrate | - | 0.44 |
|  | ml/L | ml/L |
| Tesgosept solution  (10 w/v% methyl 4-hydroxybenzoate, 0.5 v/v% 100% EtOH) | 17.46 | 7.14 |
| Propionic acid mix  (41.2 v/v% 99% propionic acid, 4.2 v/v% 85% phosphoric acid) | 9.2 | 3.57 |

*We used Bundaberg food-grade molasses, which like most molasses products, typically contains 40-55% sugar. This sugar content is mainly composed of Sucrose, Glucose and Fructose.
